# Single Cell Analysis of Regions of Interest (SCARI) using a novel photoswitchable tag

**DOI:** 10.1101/2020.10.02.291096

**Authors:** Anne M. van der Leun, Mirjam E. Hoekstra, Luuk Reinalda, Colinda L.G.J. Scheele, Mireille Toebes, Michel J. van de Graaff, Hanjie Li, Akhiad Bercovich, Yaniv Lubling, Eyal David, Daniela S. Thommen, Amos Tanay, Jacco van Rheenen, Ido Amit, Sander I. van Kasteren, Ton N. Schumacher

## Abstract

The functional activity and differentiation potential of cells is determined by their interaction with surrounding cells. Approaches that allow the unbiased characterization of cell states while at the same time providing spatial information are of major value to assess this environmental influence. However, most current techniques are hampered by a trade-off between spatial resolution and cell profiling depth. Here, we developed a photoswitch-based technology that allows the isolation and in-depth analysis of live cells from regions of interest in complex *ex vivo* systems, including human tissues. The use of a highly sensitive 4-nitrophenyl(benzofuran)-cage coupled to nanobodies allowed photoswitching of cells in areas of interest with low-intensity violet light and without detectable phototoxicity. Single cell RNA sequencing of spatially defined CD8^+^ T cells was used to exemplify the feasibility of identifying location-dependent cell states at the single cell level. Finally, we demonstrate the efficient labeling and photoswitching of cells in live primary human tumor tissue. The technology described here provides a valuable tool for the analysis of spatially defined cells in diverse biological systems, including clinical samples.

## Introduction

Methods for the in-depth characterization of individual cells, such as single cell transcriptomics and proteomics, form an essential approach for our understanding of cellular function in human tissues. While these technologies are well-suited to describe the diversity of physiological and pathophysiological cell states, information on the spatial localization of the analyzed cells is lost upon tissue dissociation. Knowledge on the spatial context of individual cells is however critical to understand how locoregional differences in environmental signals (e.g. through cell-cell interactions or soluble mediators) impact cellular state and cell differentiation. For example, within human cancers, immune cells are found both at peritumoral and intratumoral sites. In addition, intratumoral immune cells may be further subdivided into, for instance, cells located within tumor cell nests and tertiary lymphoid organs. However, the relationship between cell state and any of these different cell locations is poorly understood^1^. Classical methods that are used to simultaneously assess the localization and phenotypic properties of individual cells, such as immunohistochemistry and confocal microscopy, are limited by the number of parameters that can be analyzed. While next generation imaging techniques, such as imaging mass cytometry^2^, multi ion beam imaging by time of flight (MIBI-TOF^3^) and co-detection by indexing (CODEX^4^), allow the analysis of multiple markers on tissue slides, these technologies still require upfront decisions on the genes or proteins that are assessed, and lack the resolution offered by the former techniques.

The combination of the spatial resolution from microscopy approaches with the unbiased profiling capacity of single cell techniques has the potential to allow the *ab initio* analysis of the relationship between cell state and cellular localization. Work in transgenic mouse models has demonstrated that the localized switching of photoactivatable proteins, such as paGFP, Dronpa, or Dendra, allows the analysis of single cell transcriptomes in regions of interest with high spatial resolution^5^. However, approaches based on the genetic encoding of photoswitchable proteins are not applicable to primary human tissues. To this end, a number of methods, including Slide-seq^6^ and related spatial transcriptomics approaches^7–12^, have been developed that can be used to couple transcriptome data and spatial information from cells in human tissues. In Slide-seq based approaches, mRNA molecules from tissue slides are transferred to barcode-labeled surfaces^6–8^, thereby providing the possibility to perform an unbiased analysis of transcriptional activity at defined sites, but with the caveat that the gene expression patterns obtained by these techniques are often averages from multiple cells. The specific labeling of spatially defined cells in live tissues followed by single cell analysis of those cells upon dissociation on the other hand does not suffer from this limitation. Elegant work by the Krummel group^13^ recently demonstrated how local unshielding of DNA barcodes that are protected by photolabile 6-nitropiperonyloxylmethyl (NPOM) groups can be used to specifically mark cells in areas of interest in tissue sections. Here, we developed a distinct technology that exploits a 4-nitrophenyl(benzofuran) (NPBF) shielded protein tag to uniformly label cells *in situ* in live, intact human tissue. The use of the NPBF group allows highly-efficient uncaging under mild conditions (i.e. using low energy violet light), resulting in minimal phototoxicity. In addition, the implementation of a unique signal switch system, in which uncaging leads to the simultaneous loss of a first signal and gain of a second signal, allows the superior identification of uncaged cells. The *single cell analysis of regions of interest (*SCARI) technology that we describe can be used to specifically isolate cell populations from defined regions with high spatial resolution, and we exemplify the value of this approach through single cell RNA sequencing of human T cells at specific sites.

## Results

### Conceptual approach to achieve localized cell marking

To label cells at specific tissue sites with minimal phototoxicity, we set out to create a photoswitchable molecule in which a detectable group was shielded by a photolabile protecting group (PPG). The most commonly used PPGs for photoswitching in biological systems are the *o*-nitrobenzyl based chromophores (e.g. 6-nitroveratryloxycarbonyl (NVOC) and NPOM) that have, amongst others, been used to study T-cell activation kinetics^14^, the liberation of pro-drugs^15–18^ and variation in immune cell states^13^. These chromophores, however, have a low quantum yield (φ_u_^NVOC^ = 0.0013, φ_u_^NPOM^ = 0.0075) and therefore require either long light exposure or high energy light to remove the photolabile group, resulting in light-induced cellular damage^19^. In addition, removal of *o*-nitrobenzyl based chromophores through light exposure is accompanied by the release of toxic benzaldehyde byproducts^20–22^. The recently reported 4-nitrophenyl(benzofuran) (NPBF) chromophore^23^ on the other hand, has a superior uncaging efficiency (φ_u_^NPBF^ = 0.09) and does not release toxic byproducts upon light exposure. Furthermore, to achieve an optimal distinction between uncaged and caged cells, we designed a system in which removal of the NPBF chromophore simultaneously leads to the loss of a first detectable (fluorescent) signal and gain of a second signal. To accomplish this, we designed a FLAG-peptide (DYKDDDDK) that is protected by an Alexa Fluor 594 (AF594)-conjugated NPBF-protecting group. We envisioned that this photocage would interfere with the binding of αFLAG antibodies, and uncaging of the FLAG-tag could thus be used to simultaneously release the AF594 dye and create a novel antibody binding site (Fig. 1). To this end, a bifunctional NPBF photocleavable linker bearing an N-hydroxysuccinimide (NHS) carbonate on one end and an alkyne handle on the other end was developed to allow one to orthogonally install the photocage on the FLAG epitope and click the alkyne handle of the photocage with an azide functionalized fluorophore^24^. To be able to label specific cell types, the caged FLAG-tag was subsequently coupled to cell lineage-specific nanobodies through sortase-based reactions^25^. To this purpose, the canonical FLAG octapeptide sequence (DYKDDDDK) was extended with three N-terminal glycines^26^, to allow conjugation to LPXTG-modified nanobodies. In addition, a cysteine was placed in between the N-terminal GGG and FLAG tag sequence to prevent aspartimide formation^27^, resulting in the final sequence GGGCDYKDDDDK. The FLAG epitope contains two lysine residues that are suitable for installing the photocage. Prior work has demonstrated that the C-terminal lysine does not significantly influence antibody binding, and is thus not suitable for the intended epitope deprotection strategy^28^. On the contrary, Jungbauer and co-workers demonstrated that the N-terminal DYKD sequence of the FLAG peptide can be used in immunological detection procedures^29^, making it plausible that modification of the side chain amine of this lysine residue (Lys7) would abolish antigen recognition for at least some αFLAG antibodies.

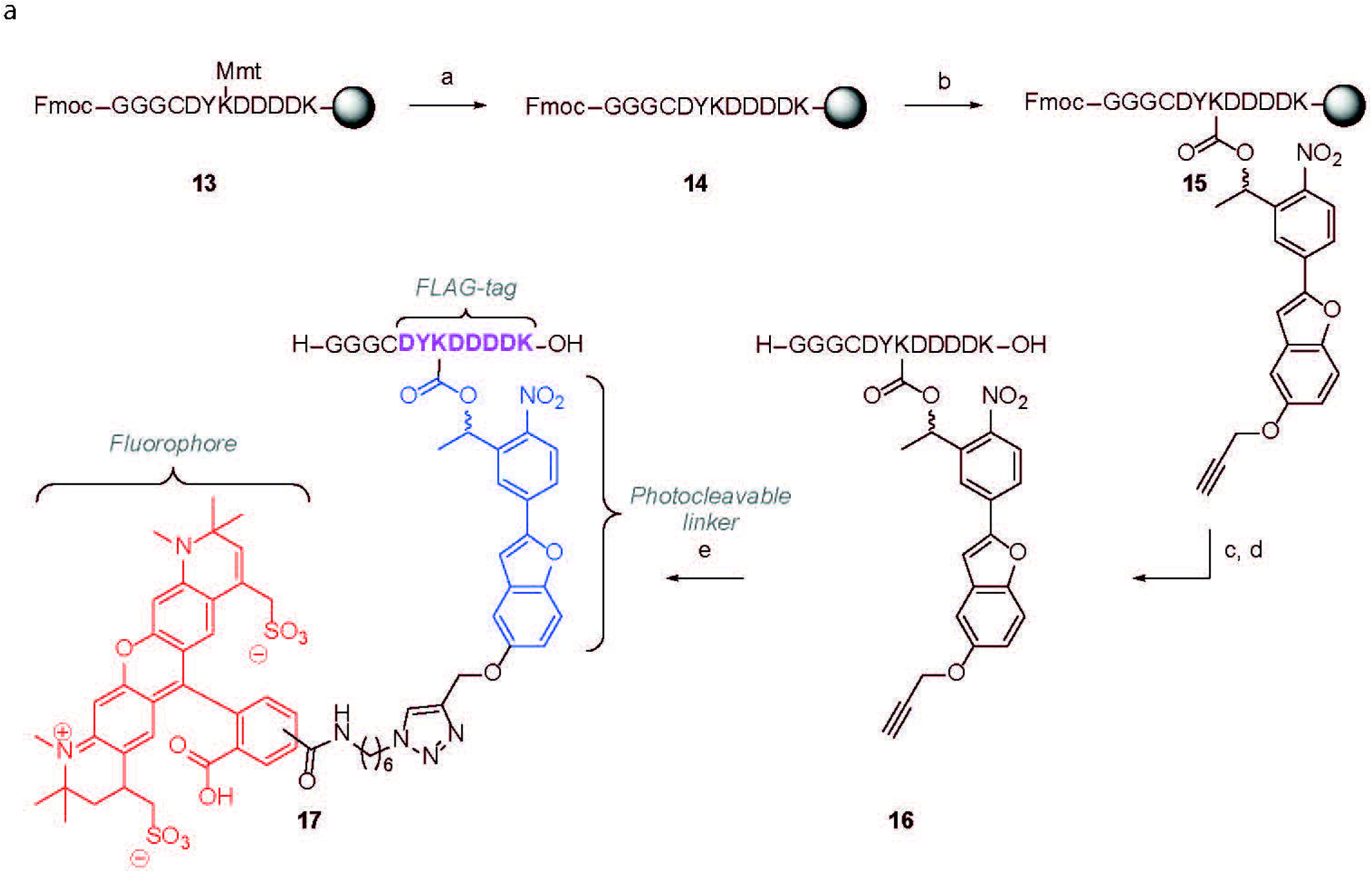

### Synthesis of the Photoswitchable Tag (PsT)

The bifunctional NPBF photocage **12**, was synthesized essentially as reported previously (Fig. S1, **1-12**)^24^. The modified FLAG tag sequence GGGCDYKDDDDK was synthesized using acid labile 4-methoxytrityl (Mmt) as the protecting group for the amine on Lys7, to allow the orthogonal deprotection of this side chain. An optimized method for Fmoc-based microwave-assisted solid-phase peptide synthesis (SPPS), which prevents racemization and aspartimide formation, was applied to obtain the core oligopeptide **13**^30^. Mmt was removed under mild conditions (1% TFA), and the sidechain amine on lys7 of **14** was reacted with the NHS carbonate of photocage **12** to yield photocaged peptide **15** on-resin. After cleavage from the resin and deprotection of all other amino acids, the photocaged peptide **16** was purified by reverse phase HPLC, followed by copper catalysed alkyne-azide cycloaddition with AF594 azide, yielding the PsT **17** (Fig. 1).

### Synthesis and characterization of photoswitchable αCD8 antibody reagents

To determine the feasibility of using PsT-labeled antibodies to selectively mark cells at defined tissue sites (Fig. 2A), we generated fluorochrome labeled camelid heavy chain-only fragments (nanobodies) specific for the human T cell marker CD8. Nanobodies display superior tissue penetration capacity as compared to regular antibodies due to their small size (∼15kDa versus ∼150kD)^31^, a property that may be particularly useful for *in vivo* or *ex vivo* staining of intact tissues with dense cellular and extracellular structures. To probe the effect of avidity on the selectivity and stability of cell marking, we first designed monomeric and dimeric fluorochrome-labeled αCD8 nanobodies. Subsequently, the stability of cell labeling when using either monomeric or dimeric αCD8 nanobodies (αCD8^M^ or αCD8^D^ respectively) was determined by staining two separate human CD8^+^ T cell populations with αCD8 nanobodies coupled to distinct fluorochromes and subsequent mixing. Following mixing of the two differentially labeled cell populations, a rapid exchange of monomeric αCD8 nanobodies was observed, resulting in the appearance of cells that were labeled with both fluorochromes. In contrast, for two different dimeric αCD8 nanobodies (αCD8^D-1^ and αCD8^D-2^) tested, highly stable cell binding was observed (Fig. 2B, Fig. S2), a property essential for the intended localized cell marking.

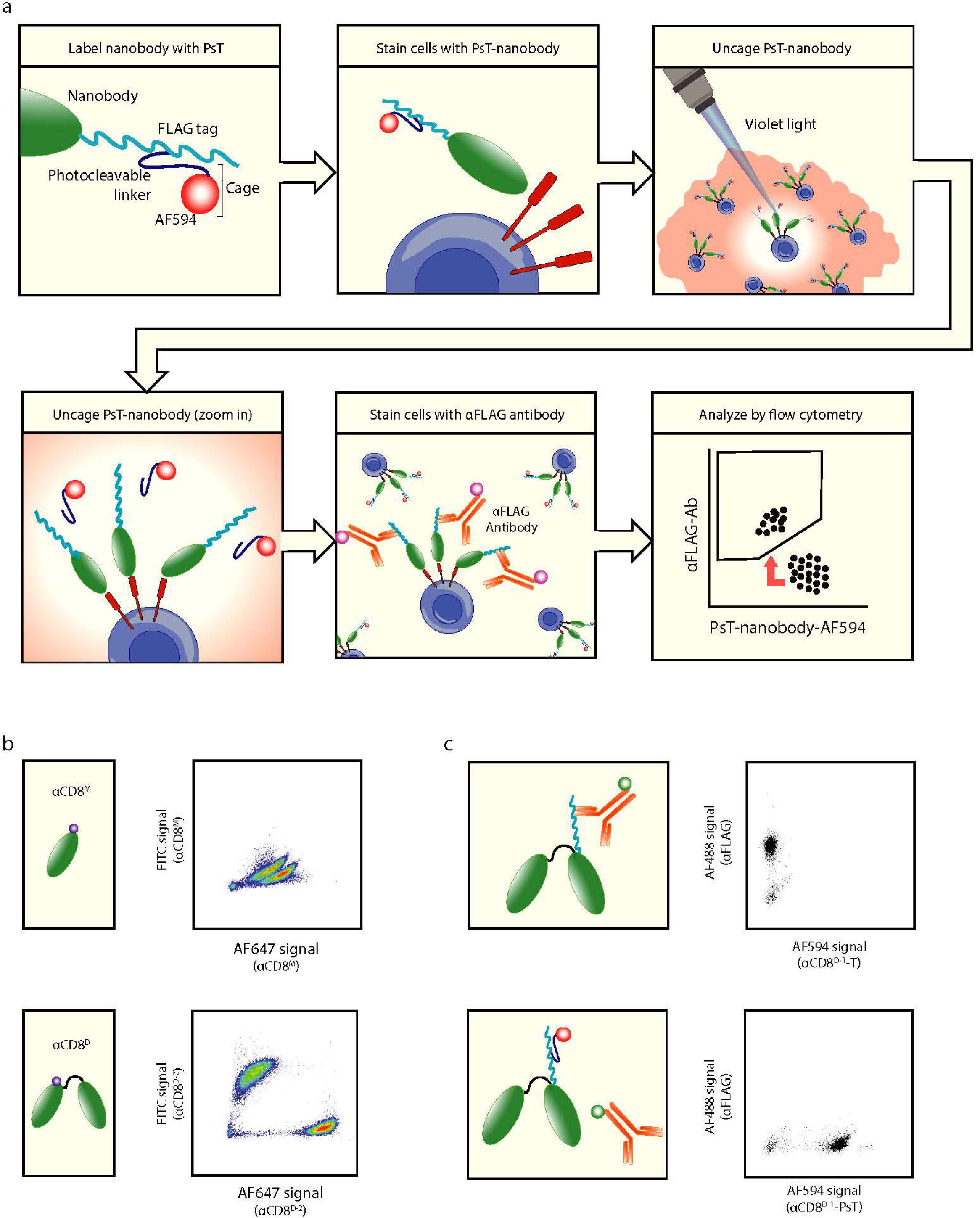

We subsequently determined whether the photoswitchable NPBF cage prevents antibody binding to the FLAG-tag. To this purpose, primary human CD8^+^ T cells were stained with FLAG-tag αCD8^D-1^ nanobodies that either contain or lack the NPBF cage (αCD8^D-1^-PsT and αCD8^D-1^-T, respectively), and accessibility of the FLAG epitope was probed using a set of different αFLAG antibodies. Notably, while certain αFLAG antibodies were insensitive to the NPBF cage (Fig. S3), binding of the D6W5B antibody was reduced to background levels upon caging of the lysine sidechain (Fig. 2C).

To understand whether photoswitching could be used to both remove the cage, and thereby the AF594 signal, and gain an αFLAG signal under conditions with limited phototoxicity, PsT labeled cells were photo-switched using a 405nm microscopy laser, and then stained with αFLAG antibody. Flow cytometric analyses of the resulting cell populations demonstrated that uncaging both led to the intended loss in AF594 signal and gain in αFLAG antibody binding (Fig. 3A-B, Fig. S4). Notably, while uncaging was already maximally effective at 865 μW/mm^2^ of violet light-exposure, cell viability remained unaffected (> 95%) up to 1440 μW/mm^2^ (Fig. 3C and Fig. S5). To explore the specificity of uncaging, we uncaged increasing surface areas of microwells containing CD8^+^ T cells in a heterogeneous population of peripheral blood mononuclear cells (PBMCs). Importantly, the fraction uncaged surface area was tightly correlated with the fraction uncaged CD8^+^ T cells, as measured by flow cytometry (Fig. 3D). Of note, mixing of cell samples that did or did not contain an uncaged CD8^+^ T cell population showed no detectable αCD8^D-1^-PsT exchange between cells, confirming the stable binding of αCD8^D-1^-PsT throughout the sample processing pipeline (Fig. 3E). Taken together, these data establish a small-protein based strategy to label and uncage cells with low light exposure, allowing isolation of specific cells by both the loss of an existing label and acquisition of a novel antibody binding site.

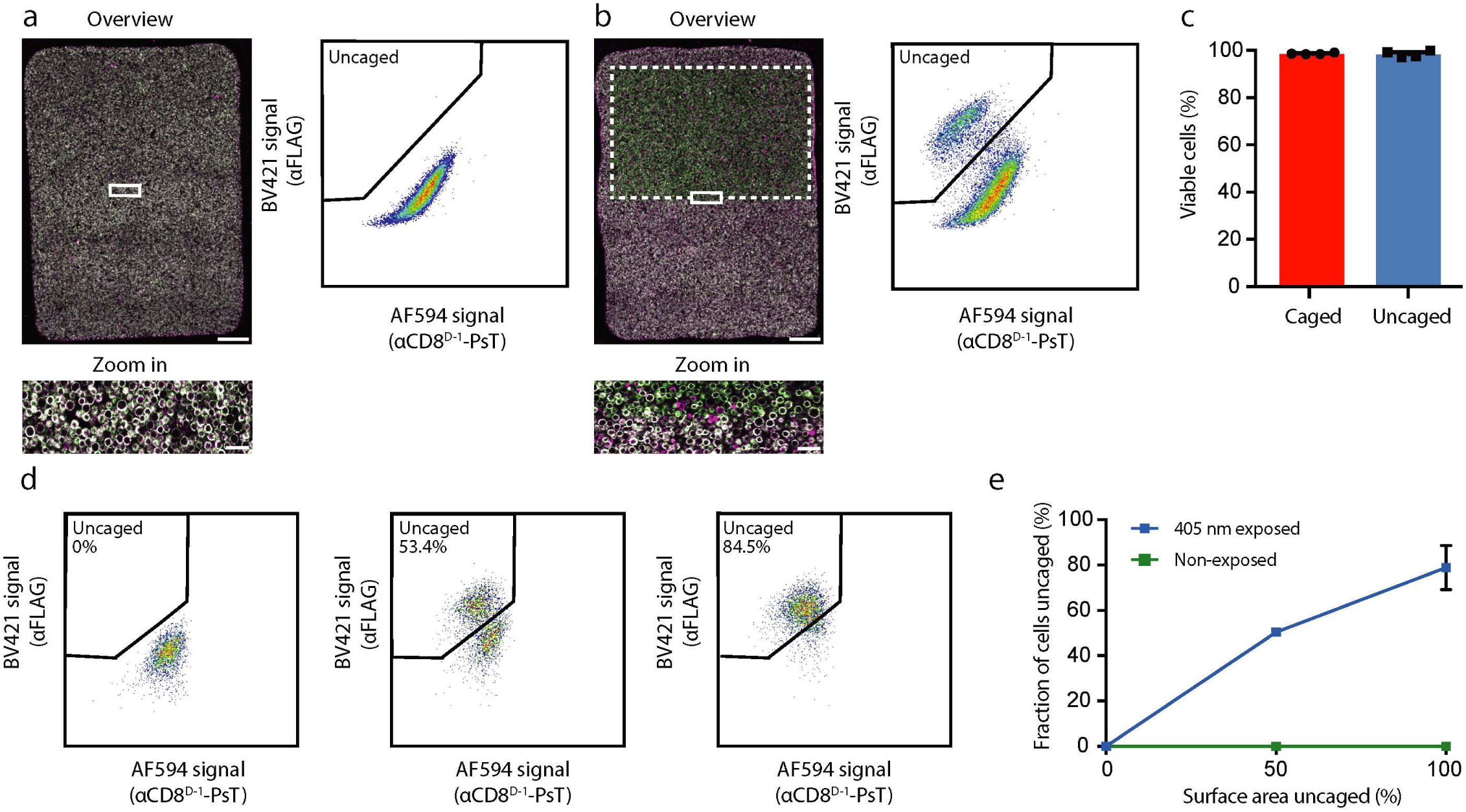

### Feasibility of local uncaging in human tumor tissue and *in vitro* cell systems

To determine the feasibility of local uncaging in more complex biological structures, we tested the efficiency of staining and uncaging of CD8^+^ T cells in viable human melanoma and non-squamous cell lung cancer (NSCLC) tissue. CD8^+^ T cells present within viable human tumor material were readily detected upon staining with αCD8^D-1^-PsT. Furthermore, uncaging of the αCD8^D-1^-PsT in melanoma and NSCLC tumors, which could be performed at single cell resolution (Fig. S6), resulted in a discrete population of AF594^low^ and αFLAG^high^ cells that was not observed in control tumor tissue (Fig. 4A and 4B), while viability of these CD8^+^ T cells remained unaffected (Fig. 4C).

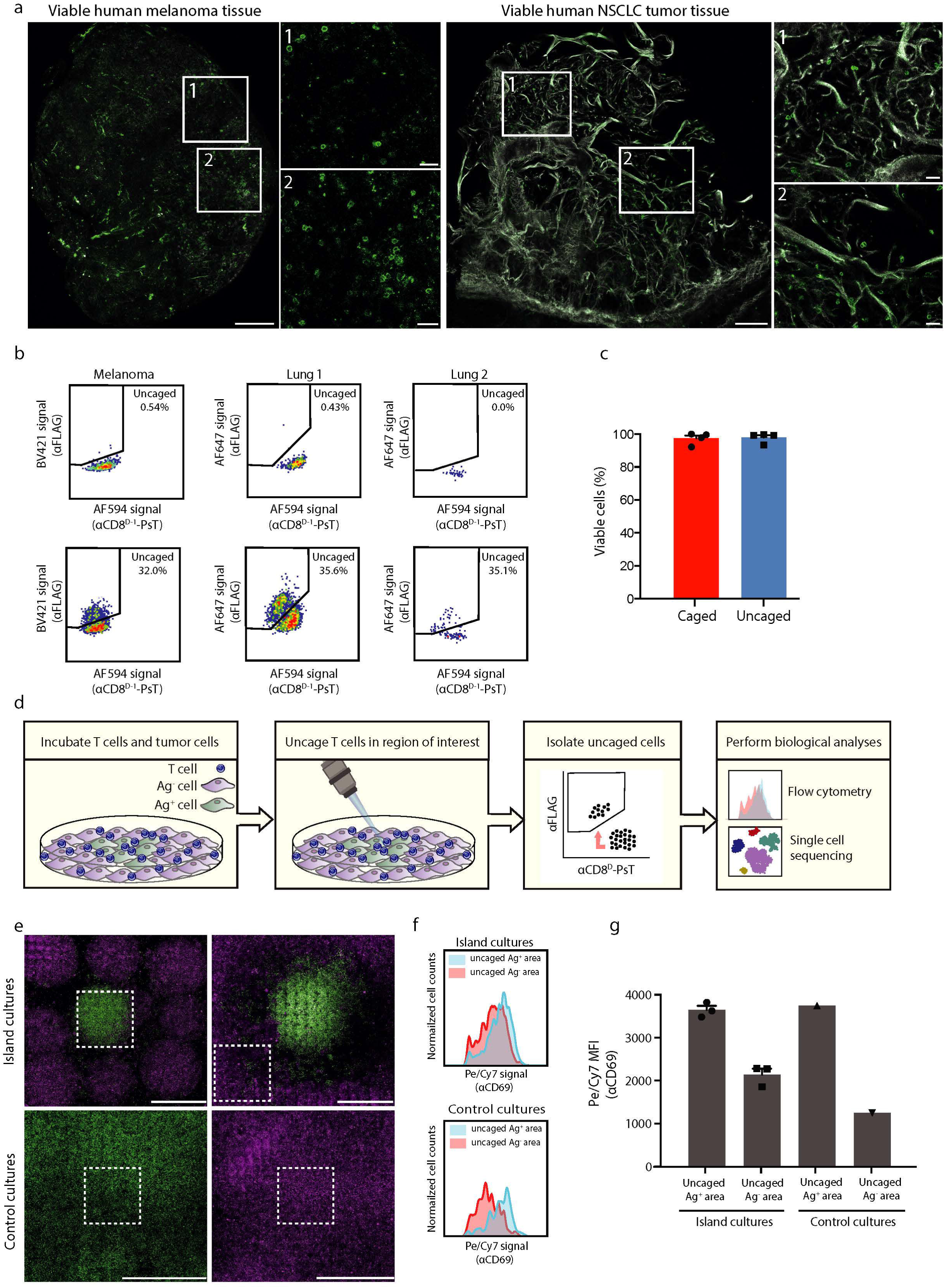

To subsequently understand whether local uncaging could be used to identify location dependent differences in cell states, we aimed to develop an *in vitro* cell system in which specific differences in cell states are induced in a controlled setting (Fig. 4D). To this purpose, adjacent islands of tumor cells that either lacked or expressed the MHC class I-restricted CDK4_R>L_ neoantigen (Katushka-labeled Ag^-^ regions and GFP-labeled Ag^+^ regions, respectively) were generated. Subsequently, αCD8^D-1^-PsT labeled CD8^+^ T cells specific for the CDK4_R>L_ neoantigen were added to such cultures, with the expectation that T cell activation would be induced in Ag^+^ but not in Ag^-^ areas. Following 4 hours of co-culture, CD8^+^ T cells in either Ag^+^ or Ag^-^ areas from island cultures were uncaged and then isolated by cell sorting (Fig. 4E). As a control, uncaged CD8^+^ T cells were isolated from separate control cultures that either only contained Ag^+^ tumor cells or Ag^-^ tumor cells. As a first test, expression of the T cell activation marker CD69 was compared on uncaged T cells that were either derived from Ag^+^ tumor cell areas or from Ag^-^ tumor cell areas. Consistent with expectations, AF594^low^ αFLAG^high^ CD8^+^ T cells isolated from cultures in which uncaging was limited to Ag^+^ tumor cell areas displayed a substantial increase in CD69 expression relative to AF594^low^ αFLAG^high^ CD8^+^ T cells from cultures in which uncaging was limited to Ag^-^ tumor cell areas (Fig. 4F-G).

### Single cell analysis of spatially defined CD8^+^ T cells

We then analyzed the transcriptomes of CD8^+^ T cells isolated from Ag^+^ and Ag^-^ regions by massive parallel single-cell RNA sequencing (MARS sequencing)^32^. To determine whether local uncaging could be used to reveal location dependent transcriptional differences, two parallel approaches were used. First, in a cell-centric approach, cell states were identified using cells from all conditions, and enrichment of specific cell states in uncaged (AF594^low^ αFLAG^high^) cells from either Ag^+^ or Ag^-^ areas was determined. Second, in gene set-centric approach, gene modules were defined based on the most variable genes in the full dataset, and differential expression of such gene modules between uncaged cells from Ag^+^ and Ag^-^ areas was subsequently analyzed.

To test for location dependent differences in cell states, T cells from all conditions were partitioned into groups of cells (“metacells”) with similar gene expression patterns, using the MetaCell algorithm^33^. This partitioning revealed a large group of T cells that unanimously expressed T cell activation markers such as TNFRSF9 and CRTAM (T-act), and a second large group of T cells that lacked expression of these marker genes (T-non-act, Fig. 5A and Fig. S7A-B, further characterization below). Notably, comparison of cell states of uncaged and caged cells in the control conditions (i.e. that only contained Ag^+^ tumor cells or only contained Ag^-^ tumor cells) demonstrated that the uncaging procedure did not influence cell states (Fig. S7C). Furthermore, uncaging did not induce detectable expression of stress-related genes (Fig. 5B-C), demonstrating that the PsT uncaging method allows for in depth analysis of viable cells, with unperturbed cell-intrinsic gene expression patterns.

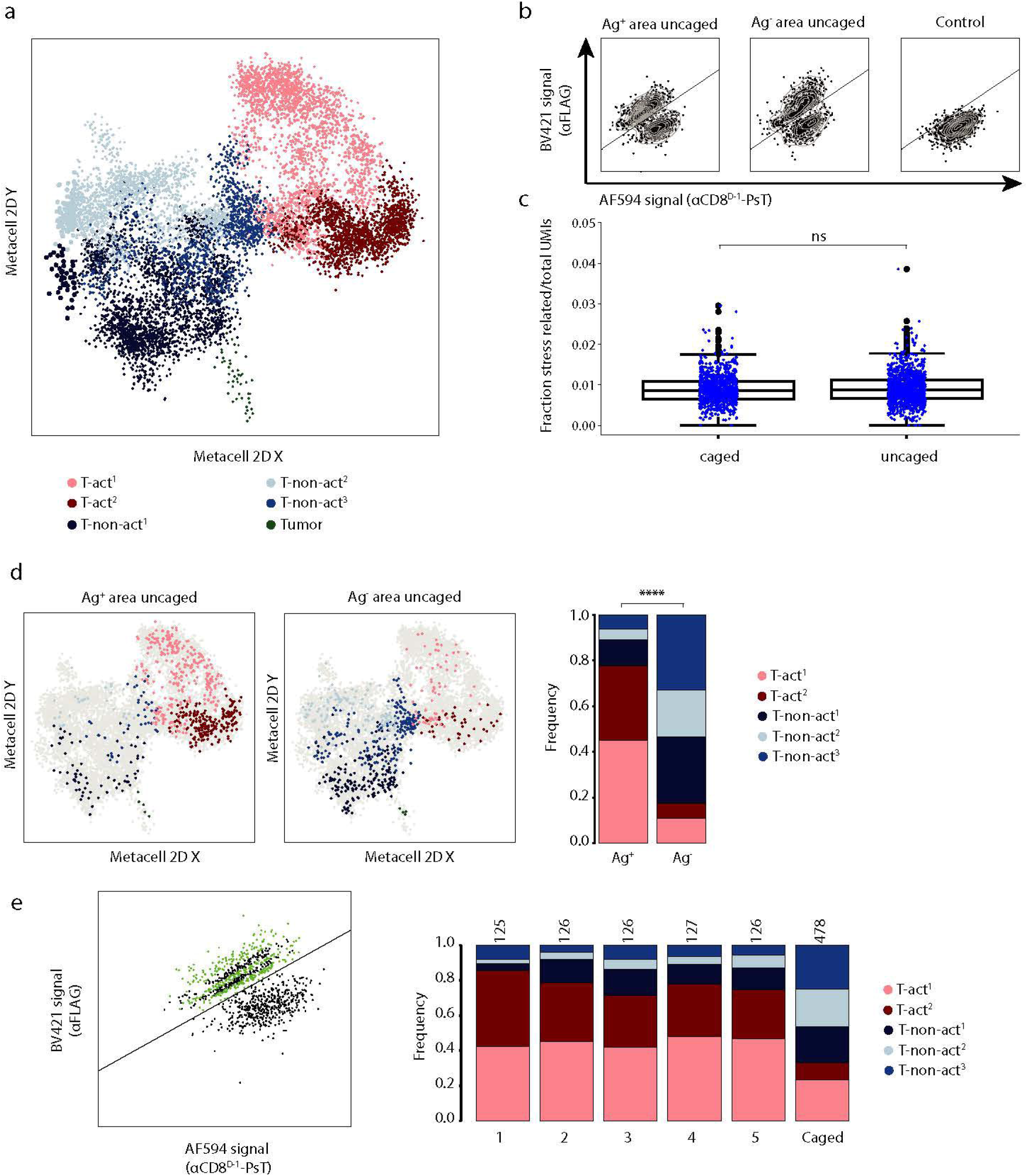

Subsequent comparison of cell states of uncaged CD8^+^ T cells derived from Ag^+^ or Ag^-^ areas revealed that the T-act state was highly enriched in Ag^+^ areas (77%), while T-non-act cells showed an increased abundance in Ag^-^ areas (82%, Fig. 5D and Fig. S7D). Also at the subgroup level (T-act^1^, T-act^2^ and T-non-act^1-3^, Fig. S7B), a substantial enrichment of activated CD8^+^ T cell states was observed in Ag^+^ areas, while all three non-activated CD8^+^ T cell populations showed enrichment in the Ag^-^ region (Fig. 5D). To determine whether the uncaged cell population is homogeneous, or whether cell pools with a lower level of uncaging (i.e. cells with intermediate AF594 and αFLAG signal) show an increased contamination with adjacent cells (with e.g. partially uncaged cells showing a non-activated cell state in samples where uncaging was aimed at Ag^+^ areas), we divided the uncaged T cells from Ag^+^ areas into bins based on their level of uncaging (bin 1 containing ‘highly uncaged’ cells to bin 5 containing ‘lowly uncaged’ cells). Notably, enrichment in activated T cell states was consistently observed across bins (Fig. 5E), demonstrating the efficient separation of cells located in different areas.

To assess whether variability in the data at the gene level could be mapped to the location of cells (i.e. in Ag^+^ or Ag^-^ areas), we next selected the top 30 genes with the highest variance throughout the entire dataset (Fig. 6A and Table S1). This list contained a considerable number of genes encoding soluble mediators, such as IFNG, CCL4, and CXCL8. Furthermore, a substantial part (67%) of the top 30 most variable genes showed increased expression in the T-act CD8^+^ T cell population (Fig. 6B). We then established a gene module containing genes with an expression pattern that was strongly correlated to that of IFNG, the most variable gene in the dataset (Fig. 6C). As a control, gene correlations to alternative anchor genes CCL4 and CXCL8 resulted in very similar gene lists (Fig. S7E). Notably, CD8^+^ T cells that were uncaged in Ag^+^ regions showed higher expression of the IFNG module than caged cells (i.e. cells from Ag^-^ areas) from the same tumor island culture (Fig. 6D). Likewise, expression of the IFNG module was significantly higher in AF594^low^ αFLAG^high^ CD8^+^ T cells that were derived from uncaged Ag^+^ regions, as compared to AF594^low^ αFLAG^high^ CD8^+^ T cells derived from uncaged Ag^-^ regions (Fig. 6E). Differential gene analysis between uncaged cells from Ag^+^ and uncaged cells from Ag^-^ areas confirmed the enrichment of soluble mediator genes as well as activation marker genes in the former cell population (Fig. S7F-G). Together, these data demonstrate that location dependent transcriptional differences can also be readily revealed at the gene level.

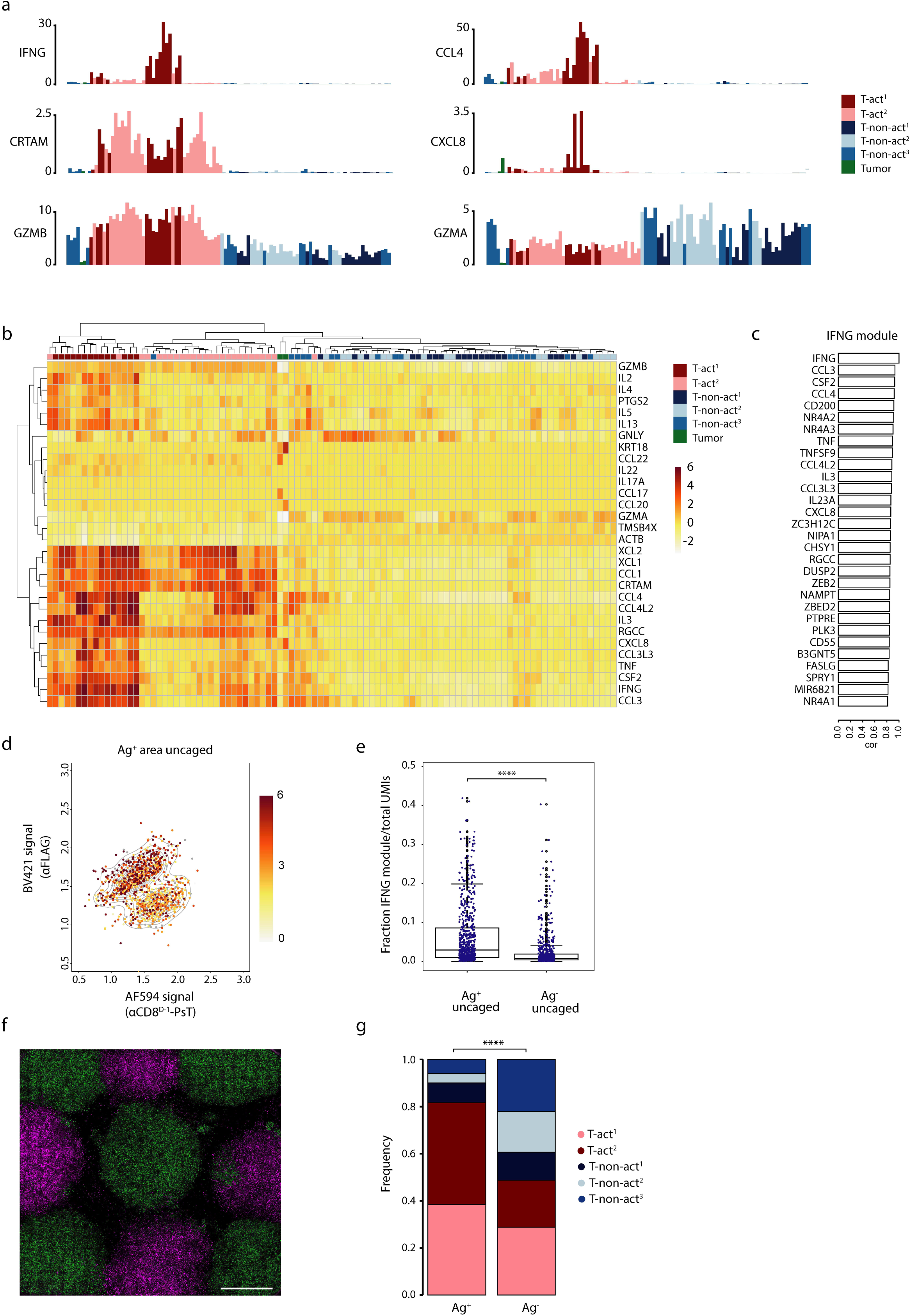

Finally, the photoswitching approach described here can also be used to identify differences in cell states in more complex systems in which many distinct adjacent cell populations are present, as tested using cultures containing a large number of intermingled areas of Ag^+^ and Ag^-^ tumor cells, with a strong enrichment of activated T cell states in Ag^+^ areas and a strong enrichment of quiescent T cell states in Ag^-^regions (Fig. 6F-G).

## Discussion

Here we present a caged protein-based technology that allows the in-depth analysis of cellular phenotypes in human tissues while preserving spatial information. The activation and differentiation state of immune cells and other cells is critically dependent on their interaction with environmental signals. For example, recent data from Ghorani et al.^34^ demonstrate that differences in the genetic make-up of distinct areas within individual human tumors coincide with variability in immune infiltrate of these areas. In other words, within human tumors, the local microenvironment is shaped by cellular interactions, either involving direct cell-cell contact or soluble mediators. To dissect how cell states are influenced by their surroundings, methods that allow in-depth analysis of cells at defined tissue sites are urgently needed. The photoswitchable tag approach presented here enables the selective isolation of cells from regions of interest in human primary tumor tissue and enables analysis of single cell transcriptomes without detectable induction of stress signatures or cell toxicity. As compared to previously developed slide-based technologies^6–12^, the method presented here has two main advantages. First, as the current approach allows isolation of viable cells from defined sites, downstream analyses are not restricted to transcriptional profiling, but can also include analysis of e.g. antigen specificity and functionality. Second, while rare cell types and cell states may be missed using grid-based approaches that do not provide full separation between neighboring cells^6–8^, the current approach does provide information that is unambiguously derived from single cell units.

The current work and the recent work by Hu et al.^13^ demonstrate how the relationship between cellular location and cell behavior can be analyzed using location dependent uncaging. The NPOM-caged DNA barcode system used in Zip-Seq^13^ is highly compatible with multiplexing and hence allows for direct comparisons between cells from multiple regions within the same tissue. On the other hand, uncaging of the NPBF chromophore as used in SCARI can be performed using low intensity violet light, rather than the more toxic ultraviolet light that is generally used to uncage photolabile groups^35^. In addition, the fact that the removal of a single NPBF chromophore suffices for uncaging in SCARI, whereas DNA barcodes in Zip-Seq are protected by four NPOM groups, plus the fact that uncaging of NPBF does not lead to release of toxic byproducts, suggests that the technique described here may be particularly suited for analysis of sensitive tissues. Finally, the sensitivity of the NPBF cage to two-photon excitation^23^ and the simultaneous gain and loss of signals that is achieved upon uncaging, will both be helpful to allow selective marking and isolation of cells located deeply into live tissues. In future work, PsT-labeled nanobodies that stably bind to pan-cell markers may, for instance, be used to identify cell types that reside in or around specific tumor structures, such as high endothelial venules or tertiary lymphoid structures^36^. Collectively, technologies such as SCARI should contribute to a further understanding on the relationship between cellular location and cell state in human tissues.

## Contributions

A.M.v.d.L., M.E.H., L.R., M.J.v.d.G., S.I.v.K., and T.N.S. conceived the project and contributed to experimental design. A.M.v.d.L., M.E.H., C.L.G.J.S. and M.T., designed, performed and analyzed biological experiments. L.R., M.T., M.J.v.d.G., and S.v.K. designed, synthesized and validated the photocaged compounds. H.L. performed single cell sequencing and E.D. performed sequence alignments. A.M.v.d.L., performed computational analysis with input from H.L., A.B., Y.L. and A.T. D.S.T. was responsible for human tumor sample acquisition. A.M.v.d.L., M.E.H., L.R., S.I.v.K., and T.N.S. wrote the manuscript with input of all other co-authors. J.v.R., I.A., S.I.v.K., and T.N.S. supervised the project.

## Acknowledgements

Plasmid sequences for αCD8 nanobodies were kindly provided by 121Bio with support of M. Gostissa and G. Grotenbreg. We thank H. Ploegh for providing the sortase expression vector. We thank M. Marqvorsen, M. de Weert, M. de Bruijn, D. Philips, D. Elatmioui, P. Hekking and staff of the NKI Flow Cytometry facility for technical support and input, and members of the van Kasteren, van Rheenen and Schumacher laboratories for discussions. This work was supported by ERC AdG SENSIT to T.N.S., and ERC StG Crosstag and ERC Cog KineTic to S.I.v.K.

## Methods

### Human material

Human tumor tissue was obtained either following opt-out procedure or upon prior informed consent, in accordance with national guidelines and after approval by the local medical ethical committee (institutional review board, IRB) of the Netherlands Cancer Institute. Tumor tissue was collected from surgical specimens after macroscopic examination of the tissue by a pathologist. Tumor tissue was dissected into fragments of 1-2mm^3^ and frozen in 90% fetal calf serum (FCS, Sigma) and 10% dimethyl sulfoxide (DMSO, Sigma). Peripheral blood mononuclear cells (PBMCs) were isolated from blood of healthy donors (Sanquin) using standard Ficoll (GE Healthcare) gradient centrifugation separation. PBMCs were stored in liquid nitrogen in 90% FCS and 10% DMSO until further use.

### General synthesis of the photocage and PsT

All reagents were commercially obtained and used without further purification unless otherwise specified. Air and moisture sensitive reagents were transferred via syringe. All air and/or moisture sensitive reactions were carried out in flame dried glassware under a positive pressure of nitrogen gas with solvents stored over activated molecular sieves (4 Å, 8-12 mesh). Reactions were monitored by analytical thin-layer chromatography (TLC) on silica gel plates (Macherey-Nagel, ALUGRAM® Xtra SIL G/UV254) and either visualized by UV light (254 nm) or by staining with a solution of KMnO_4_ (20 g/L) and K_2_CO_3_ (10 g/L) in water followed by charring at approx. 150 °C. Flash column chromatography was performed on silica gel (Macherey-Nagel, Kieselgel 60 M, 0.04 – 0.63 mm). ^1^H NMR and ^13^C-APT NMR spectra were recorded on a Bruker AV-400 (400 MHz and 101 MHz for ^1^H and ^13^C, respectively) and a Bruker AV-500 (500 MHz and 126 MHz for ^1^H and ^13^C, respectively) spectrometer at room temperature. Chemical shifts are reported in ppm relative to the residual solvent peak, the multiplicity is reported as follows: s = singlet, d = doublet, t = triplet, q = quartet, m = multiplet, br = broad signal, and J-couplings (*J*) are reported in Hertz (Hz). Liquid chromatography-mass spectrometry (LC-MS) measurements were carried out on a Finnigan Surveyor HPLC system with a Nucleodur C18 Gravity 3 µm 50 × 4.60 mm column (detection at 200 – 600 nm) coupled to a Finnigan LCQ Advantage Max mass spectrometer with electron spray ionization (ESI) or coupled to a Thermo LCQ Fleet Ion mass spectrometer with ESI. High resolution mass spectrometry (HRMS) spectra were recorded by direct injection (2 µL of a 1 µM in CH_3_CN/H_2_O/^t^BuOH) on a Thermo Scientific Q Exactive HF Orbitrap mass spectrometer equipped with an electrospray ion source in positive-ion mode (source voltage 3.5 kV, sheath gas flow 10, capillary temperature 275 °C) with resolution R = 240.000 at m/z 400 (mass range m/z 160 – 2000) correlated to an external calibration (Thermo Finnigan). Automated solid phase peptide synthesis (SPPS) was performed on a Liberty Blue™ automated microwave peptide synthesizer (CEM corporation).

### Synthesis of Bifunctional Photocage

#### 1-(2,2-diethoxyethoxy)-4-methoxybenzene (2)

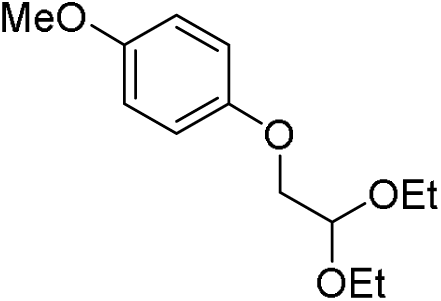

4-methoxyphenol (300 mmol, 37.2 g) was dissolved in NMP (300 mL, 1 M) followed by addition of potassium hydroxide (600 mmol, 33.6 g) at room temperature. Bromoacetaldehyde diethyl acetal (7.5 mmol, 1.13 mL) was added slowly and the reaction was stirred at 70 °C for 15 hours. The reaction mixture was poured in H_2_O and extracted with Et_2_O (3x). The combined organic layers were washed with brine (1x), dried over MgSO_4_ and concentrated *in vacuo*. The crude product was purified by flash column chromatography (pentane / Et_2_O 19:1 to 9:1) to afford compound **2** (270 mmol, 64.9 g, 90%). ^**1**^**H NMR** (400 MHz, δ (ppm), CDCl_3_): 6.90 – 6.76 (m, 4H), 4.80 (t, *J* = 5.2 Hz, 1H), 3.95 (d, *J* = 5.2 Hz, 2H), 3.80 – 3.68 (m, 2H), 3.72 (s, 3H), 3.61 (dq, *J* = 9.4, 7.0 Hz, 2H), 1.23 (t, *J* = 7.1, 6H). ^**13**^**C NMR** (101 MHz, δ (ppm), CDCl_3_): 153.9, 152.7, 115.5, 114.5, 100.5, 69.1, 62.4, 55.5, 15.3. The spectroscopic data were in accordance with those reported in our recent work^24^.

#### 5-methoxybenzofuran (3)

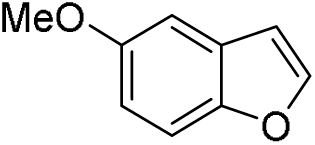

Polyphosphoric acid (60 g, H_3_PO_4_ basis, 115%) was dissolved in toluene (600 mL) and the mixture was heated to 110 °C under vigorous stirring. A solution of compound *2* (255 mmol, 61.3 g) in toluene (160 mL, 1.6 M) was added dropwise to the reaction mixture over the course of 1 hour and stirred at reflux temperature for 1 hour. The mixture was cooled to room temperature, decanted in an aqueous solution of NaOH (1 M) and extracted with toluene. The combined organic layers were washed with brine (1x), dried over MgSO_4_ and concentrated *in vacuo*. The crude product was purified by flash column chromatography (pentane / Et_2_O 100:0 to 98:2) to afford compound **3** (83.1 mmol, 12.3 g, 33%). ^**1**^**H NMR** (400 MHz, δ (ppm), CDCl_3_): 7.59 (dd, *J* = 2.2, 0.5 Hz, 1H), 7.39 (ddd, *J* = 8.9, 1.0, 0.5 Hz, 1H), 7.05 (d, *J* = 2.6 Hz, 1H), 6.90 (ddd, *J* = 8.9, 2.6, 0.5 Hz, 1H), 6.70 (dd, *J* = 2.2, 0.9 Hz, 1H), 3.84 (s, 3H). ^**13**^**C NMR** (101 MHz, δ (ppm), CDCl_3_): 145.9, 128.1, 113.2, 111.9, 106.8, 103.6, 56.03. The spectroscopic data were in accordance with those reported in our recent work^24^.

#### Benzofuran-5-ol (4)

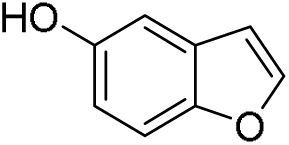

Compound *3* (5.7 mmol, 849 mg) was dissolved in anhydrous DCM (8.1 mL, 0.7 M) under N_2_ atmosphere. The mixture was cooled to -78 °C and a solution of BBr_3_ in DCM (6.8 mmol, 6.8 mL, 1 M) was slowly added and the reaction mixture was stirred at -78 °C for 1 hour and then heated to room temperature and stirred for an additional hour. The reaction was quenched with a saturated aqueous solution of NaHCO_3_, poured in H_2_O and extracted with EtOAc (3x). The combined organic layers were washed with brine (1x), dried over MgSO_4_ and concentrated *in vacuo*. The crude product was purified by flash column chromatography (pentane / EtOAc 95:5 to 9:1) to afford compound **4** (5.11 mmol, 0.685 g, 89%). ^***1***^***H NMR*** (400 MHz, δ (ppm), CDCl_3_): 7.59 (dd, *J* = 2.2, 0.5 Hz, 1H), 7.35 (ddd, *J* = 8.8, 1.0, 0.5 Hz, 1H), 7.01 (dd, *J* = 2.6, 0.6 Hz, 1H), 6.81 (ddd, *J* = 8.8, 2.6, 0.5 Hz, 1H), 6.67 (dd, *J* = 2.2, 0.9 Hz, 1H), 4.89 (s, 1H). ^**13**^**C NMR** (101 MHz, δ (ppm), CDCl_3_): 146.1, 113.1, 111.9, 106.6, 106.2. The spectroscopic data were in accordance with those reported in our recent work^24^.

#### (benzofuran-5-yloxy)(*tert*-butyl)dimethylsilane (5)

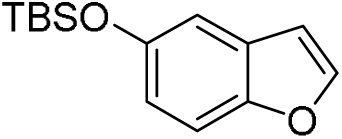

Compound **15** (3.73 mmol, 0.501 g) was dissolved in anhydrous DMF (3.8 mL, 1 M) under N_2_ atmosphere. Imidazole (18.7 mmol, 1.27 g) and a 50 wt% solution of *tert*-butyldimethylsilyl chloride in toluene (11.3 mmol, 3.9 mL, 2.9 M) were added and the reaction mixture was stirred for 1 hour at room temperature. The mixture was then poured in H_2_O and extracted with Et_2_O (3x). The combined organic layers were washed with brine (1x), dried over MgSO_4_ and concentrated *in vacuo*. The crude product was purified by flash column chromatography (pentane / Et_2_O 99:1) to afford compound *5* (2.62 mmol, 0.65 g, 70.2%). ^**1**^**H NMR** (400 MHz, δ (ppm), CDCl_3_): 7.60 (d, *J* = 2.1 Hz, 1H), 7.38 (dd, *J* = 8.8, 0.9 Hz, 1H), 7.06 (d, *J* = 2.5 Hz, 1H), 6.85 (dd, *J* = 8.8, 2.4 Hz, 1H), 6.69 (dd, *J* = 2.2, 0.9 Hz, 1H), 1.05 (d, *J* = 0.9 Hz, 9H), 0.24 (d, *J* = 0.8 Hz, 6H). ^**13**^**C NMR** (101 MHz, δ (ppm), CDCl_3_): 151.5, 150.5, 145.8, 128.2, 117.6, 111.6, 111.1, 106.7, 30.5, 25.9, 18.4, -4.3. The spectroscopic data were in accordance with those reported in our recent work^24^.

#### (5-((*tert*-butyldimethylsilyl)oxy)benzofuran-2-yl)boronic acid (6)

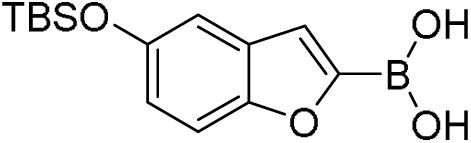

Compound **6** (3.65 mmol, 0.906 g) was dissolved in anhydrous THF (20 mL, 0.2 M) under N_2_ atmosphere and cooled to -78 °C. A solution of n-butyllithium in hexanes (8.0 mmol, 5.02 mL, 1.6 M) was added dropwise and the mixture was stirred for 1 hour. Triisopropyl borate (4.4 mmol, 1.0 mL) was added dropwise and the mixture was stirred for another 30 minutes and then heated to room temperature and stirred for an additional 30 minutes. The reaction mixture was quenched with a solution of HCl (aq.) (4 mL, 8 mmol, 2 M), poured in H_2_O and extracted with EtOAc (3x). The combined organic layers were washed with H_2_O (1x) and brine (1x), dried over MgSO_4_ and concentrated *in vacuo*. The crude product **6** was used as is in the following reaction.

#### 5’-bromo-2’-nitroacetophenone (7)

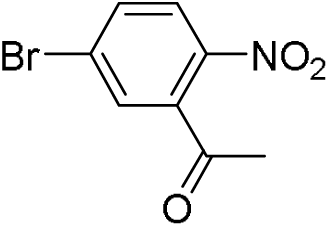

KNO_3_ (24.0 mmol, 2.42 g) was loaded into a flask and cooled down to -20°C. H_2_SO_4_ (21.75 mL, 98%) was added and the mixture was stirred for 30 minutes. 3’-bromoacetophenone (20 mmol, 3.98 g, 2.64 mL) was slowly added. The reaction mixture was heated up to -10 °C and stirred for 2 hours. The reaction mixture was poured into crushed ice and extracted with DCM (3x). The combined organic layers were washed with brine (1x), dried over MgSO_4_ and concentrated *in vacuo*. The crude product was recrystallized from Et_2_O / pentane yielding compound **7** (9.8 mmol, 2.4 g, 49%). ^**1**^**H NMR** (400 MHz, δ (ppm), CDCl_3_): 8.00 (d, *J* = 8.7 Hz, 1H), 7.74 (dd, *J* = 8.7, 2.1 Hz, 1H), 7.55 (d, *J* = 2.1 Hz, 1H), 2.56 (s, 3H). ^**13**^**C NMR** (101 MHz, δ (ppm), CDCl_3_): 133.8, 130.5, 126.1, 30.4. The spectroscopic data were in accordance with those reported in our recent work^24^.

#### 1-(5-(5-((tert-butyldimethylsilyl)oxy)benzofuran-2-yl)-2-nitrophenyl)ethan-1-one (8)

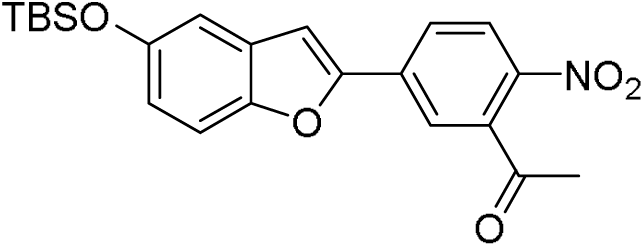

Compound *6* (4.62 mmol, 1.35 g) and compound *7* (3.65 mmol, 0.89 g) were dissolved in a 1:1 mixture of THF and H_2_O (42 mL) under N_2_ atmosphere. The flask was wrapped in aluminum foil and K_2_CO_3_ (5.48 mmol, 0.757 g) and Pd(Ph_3_)_4_ (0.23 mmol, 0.266 g) were added. The mixture was heated to 66 °C and refluxed for 18 hours. The reaction mixture was quenched with a saturated aqueous solution of NH_4_Cl, poured in H_2_O and extracted with EtOAc (3x). The combined organic layers were washed with brine (1x), dried over MgSO_4_ and concentrated *in vacuo*. The crude product was purified by flash column chromatography (pentane / Et_2_O 19:1 to 7:3) to afford compound **8** (2.23 mmol, 0.919 g, 61%). Increasing the polarity of the eluent (pentane / EtOAc 7:3) yielded desilylated compound **9** (0.51 mmol, 0.15 g, 14%). ^**1**^**H NMR** (400 MHz, δ (ppm), CDCl_3_): 8.09 (d, *J* = 8.6 Hz, 1H), 7.87 (dd, *J* = 8.6, 1.9 Hz, 1H), 7.76 (d, *J* = 1.8 Hz, 1H), 7.36 (dt, *J* = 8.9, 0.8 Hz, 1H), 7.13 (d, *J* = 0.9 Hz, 1H), 7.05 (d, *J* = 2.4 Hz, 1H), 6.89 (dd, *J* = 8.8, 2.5 Hz, 1H), 2.61 (s, 3H), 1.04 (d, *J* = 0.7 Hz, 13H), 0.25 (d, *J* = 0.7 Hz, 7H). ^**13**^**C NMR** (101 MHz, δ (ppm), CDCl_3_): 199.76, 152.82, 152.03, 150.87, 144.03, 139.06, 136.13, 129.17, 125.65, 125.14, 122.72, 119.69, 111.72, 111.24, 105.94, 30.24, 25.69, 18.18, -4.48. The spectroscopic data were in accordance with those reported in our recent work^24^.

#### 1-(5-(5-hydroxybenzofuran-2-yl)-2-nitrophenyl)ethan-1-one (9)

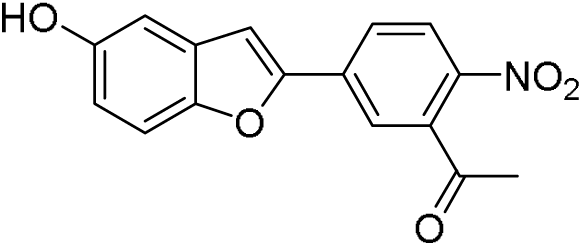

Compound **8** (0.66 mmol, 0.27 g) was dissolved in anhydrous THF (7 mL) under N_2_ atmosphere. A solution of 70% HF in Pyridine (0.7 mL, 10%) was added and the reaction was stirred at room temperature for 2 hours. The reaction was quenched with a saturated aqueous solution of NaHCO_3_ until no CO_2_ release was observed. The mixture was poured into H2O and extracted with EtOAc (3x). The combined organic layers were washed with brine (1x), dried over MgSO_4_ and concentrated *in vacuo*. The crude product was purified by flash column chromatography (pentane / EtOAc 17:3 to 7:3) to afford compound **9** (0.62 mmol, 0.18 g, 93%). ^**1**^**H NMR** (400 MHz, δ (ppm), DMSO-*d6*): 9.40 (s, 1H), 8.22 (d, *J* = 8.4 Hz, 1H), 8.17 – 8.09 (m, 2H), 7.67 (s, 1H), 7.47 (d, *J* = 8.8 Hz, 1H), 7.01 (d, *J* = 2.5 Hz, 1H), 6.86 (dd, *J* = 8.9, 2.5 Hz, 1H), 2.63 (s, 3H). ^**13**^**C NMR** (101 MHz, δ (ppm), DMSO-*d6*): 199.8, 154.0, 152.6, 149.3, 144.3, 138.0, 135.3, 129.2, 126.2, 125.6, 123.2, 115.4, 111.9, 106.8, 105.8, 30.1. The spectroscopic data were in accordance with those reported in our recent work^24^.

#### 1-(2-nitro-5-(5-(prop-2-yn-1-yloxy)benzofuran-2-yl)phenyl)ethan-1-one (10)

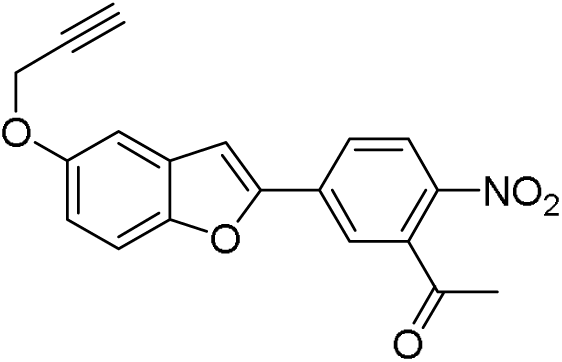

Compound *9* (0.44 mmol, 127 mg) was dissolved in anhydrous DMF (3.5 mL). K_2_CO_3_ (1.8 mmol, 0.25 g) was added and the reaction was stirred at room temperature for 1 hour under N_2_ atmosphere. Then, propargyl bromide (0.90 mmol, 0.10 mL) was added dropwise and the resulting suspension was stirred for 5 hours. The reaction mixture was quenched with a saturated aqueous solution of NH_4_Cl, extracted with EtOAc (3 × 15 mL). The combined organic layers were washed with brine (50 mL), dried over MgSO_4_ and concentrated *in vacuo*. The crude product was purified by flash column chromatography (pentane / DCM 2:1 to 3:1) to afford compound **10** (0.38 mmol, 0.13 g, 86%). ^**1**^**H NMR** (400 MHz, δ (ppm), Acetone-*d6*): 8.25 (dd, *J* = 8.6, 0.5 Hz, 1H), 8.21 (dd, *J* = 8.6, 1.9 Hz, 1H), 8.13 (dd, *J* = 1.9, 0.5 Hz, 1H), 7.65 (d, *J* = 0.9 Hz, 1H), 7.57 (ddd, *J* = 8.9, 0.9, 0.5 Hz, 1H), 7.33 (dd, *J* = 2.6, 0.5 Hz, 1H), 7.09 (dd, *J* = 8.9, 2.6 Hz, 1H), 4.85 (d, *J* = 2.4 Hz, 2H), 3.10 (t, *J* = 2.4 Hz, 1H), 2.64 (s, 3H). ^**13**^**C NMR** (101 MHz, δ (ppm), Acetone-*d6*): 199.8, 155.6, 154.4, 151.8, 145.9, 139.7, 136.6, 130.3, 127.0, 126.3, 124.2, 116.8, 112.9, 107.3, 106.5, 79.8, 77.1, 57.1, 30.2. **HRMS:** Calculated for C_19_H_14_NO_5_ 336.08665 [M+H]^+^; Found 336.08664

#### 1-(2-nitro-5-(5-(prop-2-yn-1-yloxy)benzofuran-2-yl)phenyl)ethan-1-ol (11)

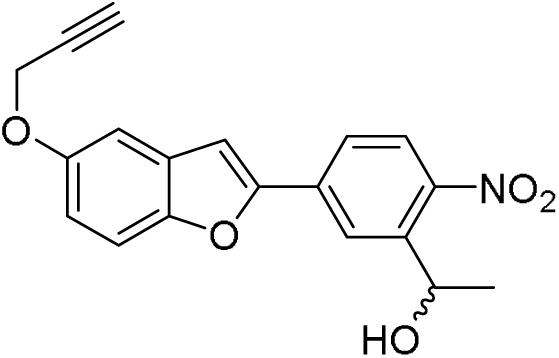

Compound **10** (0.30 mmol, 0.10 g) was dissolved in 1,4-dioxane (12 mL), then diluted with MeOH (9 mL) and the solution was cooled to 0 °C. NaBH4 (0.45 mmol, 17 mg) was added portionwise and the reaction mixture was stirred at room temperature for 1 hour. The reaction was quenched with acetone (10 mL) and the volatile solvents were removed by a gentle stream of N_2_. The reaction mixture was diluted with EtOAc (25 mL), washed with water (2 × 25 mL) and brine (25 mL) and the organic layer was dried over MgSO_4_ and concentrated *in vacuo*. The crude product was purified by flash column chromatography (pentane / EtOAc 1:9 to 1:4) to afford compound **11** (0.26 mmol, 88 mg, 87%). ^**1**^**H NMR** (400 MHz, δ (ppm), Acetone-*d6*): 8.46 (d, *J* = 1.8 Hz, 1H), 8.04 (d, *J* = 8.3 Hz, 1H), 7.99 (dd, *J* = 8.5, 1.9 Hz, 1H), 7.57 (dt, *J* = 8.9, 0.7 Hz, 1H), 7.54 (d, *J* = 0.9 Hz, 1H), 7.31 (dd, *J* = 2.6, 0.5 Hz, 1H), 7.06 (dd, *J* = 8.9, 2.6 Hz, 1H), 5.47 (dtd, *J* = 6.7, 6.0, 4.2 Hz, 1H), 4.85 (d, *J* = 2.4 Hz, 2H), 4.74 (d, *J* = 4.1 Hz, 1H), 3.11 (t, *J* = 2.4 Hz, 1H), 1.54 (d, *J* = 6.3 Hz, 3H). ^**13**^**C NMR** (101 MHz, δ (ppm), Acetone-*d6*): 155.3, 151.5, 147.5, 144.5, 135.6, 130.3, 125.8, 124.5, 116.1, 112.7, 106.2, 105.9, 79.8, 77.0, 65.6, 65.5, 57.0, 25.5, 25.4. The spectroscopic data were in accordance with those reported in our recent work^24^.

#### 2,5-dioxopyrrolidin-1-yl (1-(2-nitro-5-(5-(prop-2-yn-1-yloxy)benzofuran-2-yl)phenyl)ethyl) carbonate (12)

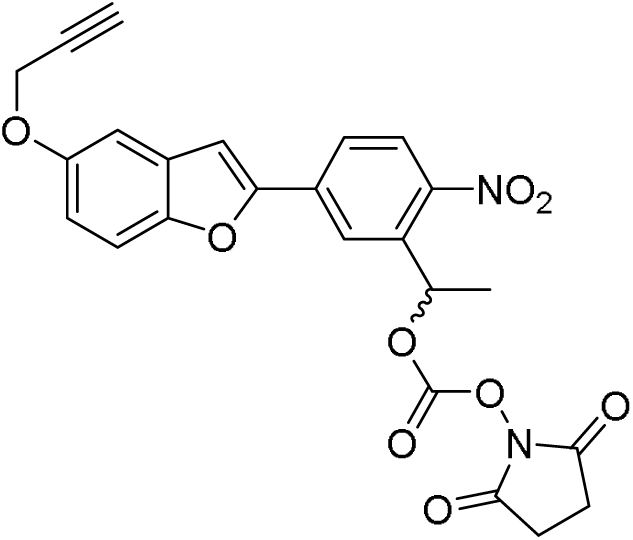

Compound **11** (0.19 mmol, 63 mg) and disuccinimidyl carbonate (0.37 mmol, 96 mg) were dissolved in anhydrous DMF (2 mL) at room temperature under N_2_ atmosphere. Et_3_N (0.56 mmol, 78 µL) was added slowly and the reaction mixture was stirred for 4 hours. The resulting solution was diluted with EtOAc (50 mL), washed with water (3 × 50 mL) and brine (50 mL), dried over MgSO_4_ and concentrated *in vacuo*. The crude product was purified by flash column chromatography (pentane / EtOAc 4:1 to 2:1) to afford compound **12** (0.15 mmol, 71 mg, 79%). ^**1**^**H NMR** (400 MHz, δ (ppm), DMSO-*d6*): 8.21 (d, *J* = 1.9 Hz, 1H), 8.19 (d, *J* = 8.6 Hz, 1H), 8.13 (dd, *J* = 8.6, 1.9 Hz, 1H), 7.80 (d, *J* = 0.9 Hz, 1H), 7.65 (dt, *J* = 9.1, 0.7 Hz, 1H), 7.32 (d, *J* = 2.6 Hz, 1H), 7.06 (dd, *J* = 9.0, 2.6 Hz, 1H), 6.34 (q, *J* = 6.4 Hz, 1H), 4.86 (d, *J* = 2.4 Hz, 2H), 3.59 (t, *J* = 2.4 Hz, 1H), 2.77 (s, 4H), 1.81 (d, *J* = 6.5 Hz, 3H). ^**13**^**C NMR** (101 MHz, δ (ppm), Acetone-*d6*): 169.8, 154.0, 152.8, 150.5, 150.2, 146.4, 136.0, 135.1, 128.6, 126.0, 125.4, 122.7, 115.7, 112.2, 106.6, 105.6, 79.5, 78.3, 75.4, 56.1, 25.4, 21.1. *HRMS:* Calculated for C_24_H_18_N_2_NaO_9_ 501.09045 [M+Na]^+^; Found 501.09049

### Synthesis of Photoswitchable Tag

#### Fmoc-GGGCDYK(Mmt)DDDDK on resin (13)

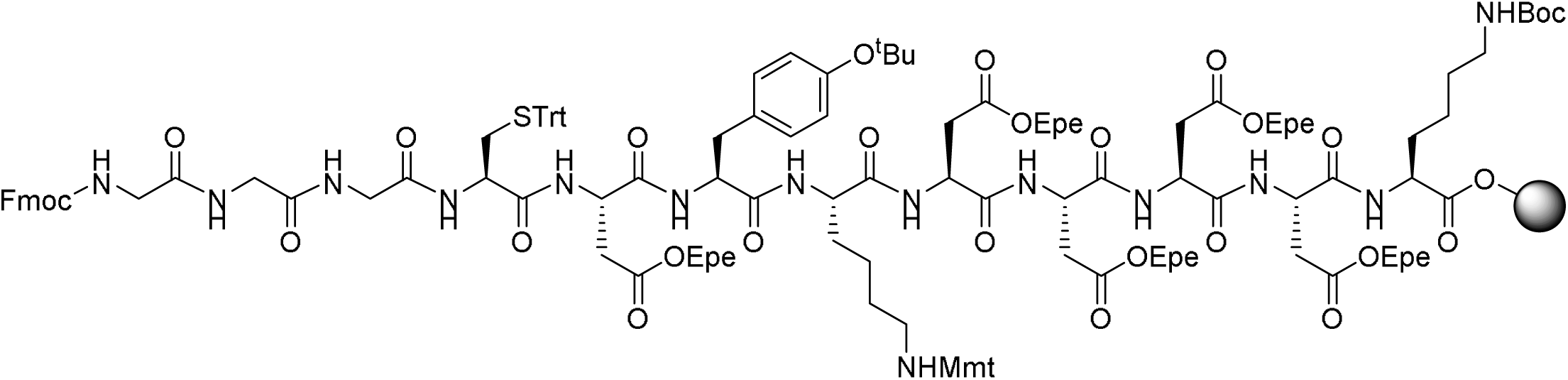

The peptide sequence was synthesized using Fmoc SPPS starting from Tentagel™ S PHB-Lys(Boc)Fmoc (100 µmol). Fmoc protected amino acids (AA) Fmoc-Asp(OEpe)-OH, Fmoc-Lys(Mmt)-OH, Fmoc-Tyr(*t*Bu)-OH, Fmoc-Cys(Trt)-OH and Fmoc-Gly-OH were coupled (order from right to left: GGGCDYKDDDD) with the following automated and repetitive sequence: 1) Fmoc deprotection with 20% piperidine and 1% formic acid in DMF at room temperature (1^st^ cycle 5 minutes, 2^nd^ cycle 10 minutes). 2) coupling with AA (5 eq.) and equimolar ethyl 2-cyano-2-(hydroxyamino)acetate (OxymaPure®) (1.0 M in DMF, with 0.4 M DiPEA in the same solution) and *N,N’*-diisopropylcarbodiimide (DIC) (0.5 M in DMF) at 90 °C for 2.5 minutes (microwave, 30 W). The resin was washed with DMF (5 × 5 mL) and a small sample was taken for LC-MS analysis. The sample (approx. 10 mg) was deprotected (acid labile) and cleaved from the resin with 0.5 mL TFA/TIS/H_2_O (95/2.5/2.5) on the shaker for 3 h. The solution was precipitated in cold Et_2_O (2.5 mL), centrifuged for 3 minutes at 4500 rpm and the supernatant removed. The pellet was dissolved in 3 mL CH_3_CN/H_2_O/^t^BuOH. **LC-MS** (linear gradient 30 – 70% MeCN, 0.1% TFA, 13 minutes) R_t_ (minutes): 1.61, ESI (m/z): 1509.5 [M+H]^+^ for deprotected and cleaved Fmoc-GGGCDYKDDDDK-OH

#### Mmt deprotected Fmoc-GGGCDYKDDDDK on resin (14)

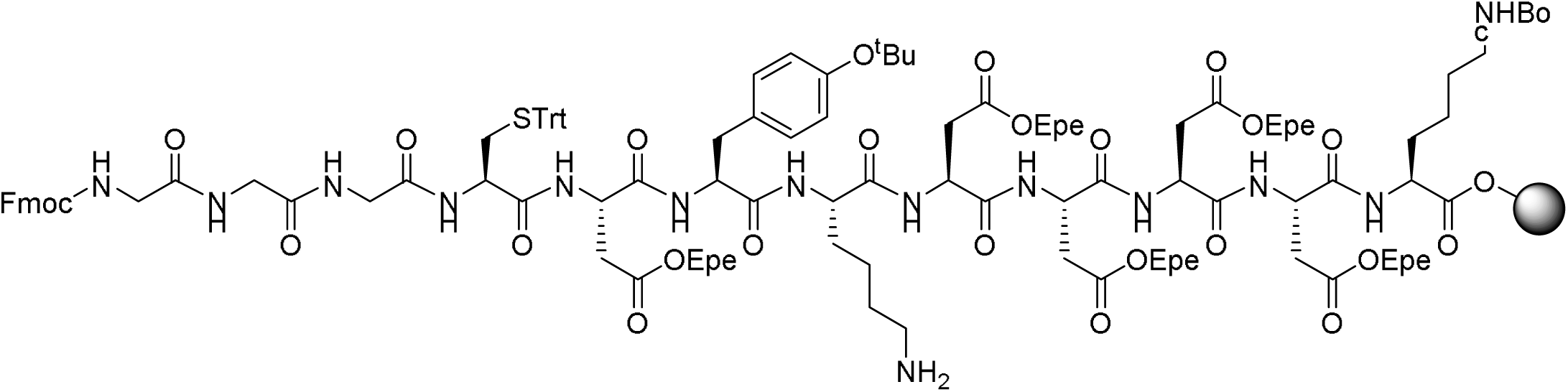

Resin **13** (90 µmol) was dissolved in 1% v/v TFA in DCM under gentle agitation at room temperature (10 × 2 minutes). A small sample (approx. 10 mg) was acetylated with 1 mL DMF/Ac_2_O/DiPEA (85/10/5) (2 × 2 minutes), washed with DMF (5 × 3 mL) and taken for LC-MS analysis. The sample (approx. 10 mg) was deprotected (acid labile) and cleaved from the resin with 0.5 mL TFA/TIS/H_2_O (95/2.5/2.5) under gentle agitation for 3 hours. The solution was precipitated in cold Et_2_O (2.5 mL), centrifuged for 3 minutes at 4500 rpm and the supernatant removed. The pellet was dissolved in 3 mL CH_3_CN/H_2_O/^t^BuOH. **LC-MS** (linear gradient 30 – 70% MeCN, 0.1% TFA, 13 minutes) R_t_ (minutes): 2.19, ESI (m/z): 1552.3 [M+1+H]^+^ for deprotected, cleaved and acetylated Fmoc-GGGCDYK(Ac)DDDDK-OH

#### Photocaged Fmoc-GGGCDYK(PC)DDDDK on resin (15)

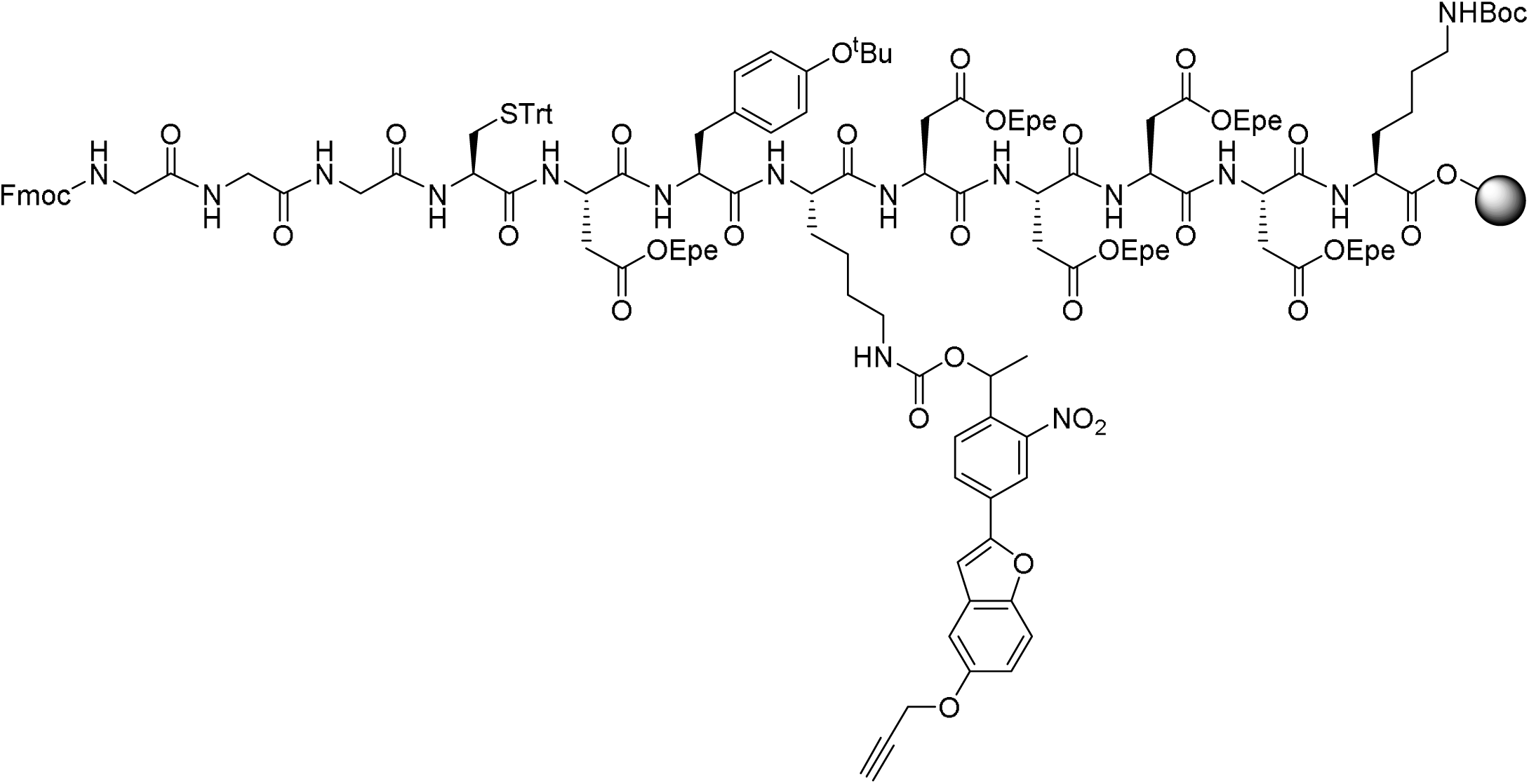

A solution of compound **12** (120 µmol, 58 mg) in DMF (1 mL) and DiPEA (75 µmol, 21 µL) were added to resin **14** (80 µmol) and shaken for 18 hours at room temperature. The resin was washed with DMF (5 × 2 mL) and a small sample was taken for LCMS analysis. The sample (approx. 10 mg) was deprotected (acid labile) and cleaved from the resin with 0.5 mL TFA/TIS/H_2_O (95/2.5/2.5) under gentle agitation for 3 hours. The solution was precipitated in cold Et_2_O (2.5 mL), centrifuged for 3 minutes at 4500 rpm and the supernatant removed. **LC-MS** (linear gradient 30 – 70% MeCN, 0.1% TFA, 13 minutes) R_t_ (minutes): 5.78, ESI (m/z): 1873.5 [M+1+H]^+^ for deprotected and cleaved Fmoc-GGGCDYK(PC)DDDDK-OH

#### Photocaged H-GGGCDYKDDDDK-OH (16)

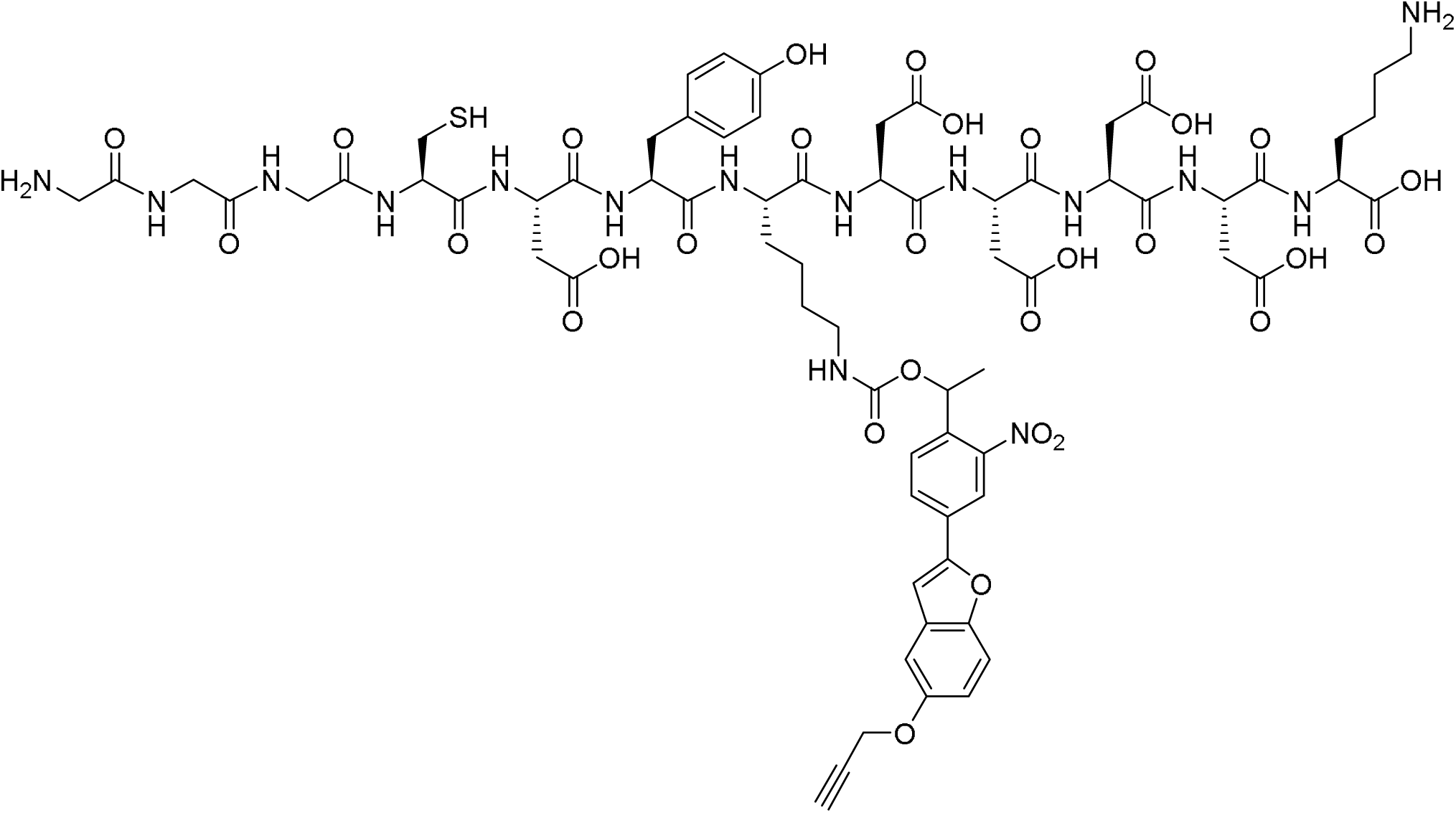

Resin **15** (70 µmol) was dissolved in 20% v/v piperidine in DMF (1 mL) under gentle agitation at room temperature (1^st^ and 2^nd^ cycle 2 minutes, 3^rd^ cycle 5 minutes). The resin was washed with DMF (5 × 2 mL) and shaken in 1 mL TFA/TIS/H_2_O (95/2.5/2.5) for 3 hours. The solution was precipitated in cold Et_2_O (5 mL), centrifuged for 3 minutes at 4500 rpm and the supernatant removed. The crude peptide was purified by RP-HPLC. **LC-MS** (linear gradient 30 – 70% MeCN, 0.1% TFA, 11 minutes) R_t_ (minutes): 4.23, ESI (m/z): 1650.5 (M + H^+^) for deprotected and cleaved H-GGGCDYK(PC)DDDDK-OH. **HRMS:** Calculated for C_70_H_89_N_15_O_30_S 1651.56095 [M+H]^2+^; Found 1651.56094

#### Photocaged AF594 H-GGGCDYKDDDDK-OH (17)

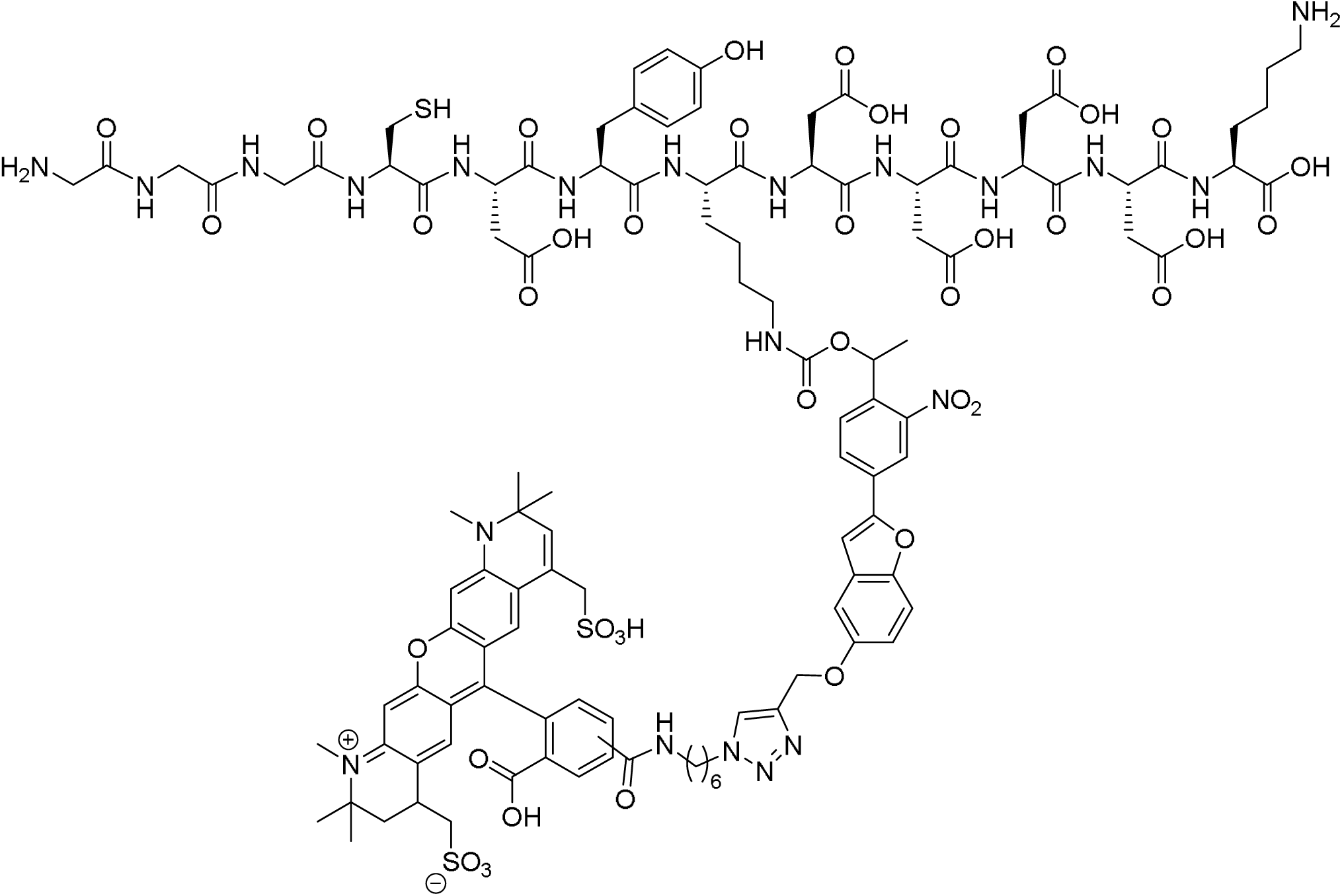

Compound **17** (0.49 µmol, 0.81 mg) was dissolved in a solution of AFDye™ 594 Azide in DMSO (0.01 M, 0.44 µmol, 44 µL) in a dark vial and diluted with ^t^BuOH (106 µL) and H_2_O (27 µL). An aqueous solution of CuSO_4_ ·5 H_2_O (0.01 M, 0.49 µmol, 49 µL), THPTA (0.01 M, 0.25 µmol, 25 µL) and (+)-sodium L-ascorbate (0.1 M, 4.9 µmol, 49 µL) were mixed, added to the dark vial and stirred at room temperature under N_2_ atmosphere for 3 hours. The reaction was quenched with an aqueous solution of EDTA (0.5 M, 5.0 µmol, 10 µL) and the reaction mixture was precipitated in ice cold milliQ (10 mL), centrifuged for 3 minutes at 4500 rpm and the supernatant removed. **LC-MS** (linear gradient 30 – 70% MeCN, 0.1% TFA, 11 minutes) R_t_ (minutes): 2.77, ESI (m/z): 1249.8 [M+H]^2+^ extracted from crude reaction mixture.

### Production of recombinant nanobodies in Escherichia coli

Monomeric and dimeric variants of two αCD8 nanobody clones (αCD8^M^, αCD8^D-1^, αCD8^D-2^) were generated as follows: *Escherichia coli* WK6 cells were transformed with the pHEN6 expression vector (for production of the αCD8^M^ nanobody) and *Escherichia coli* BL21 cells were transformed with the pET22b expression vector (for production of the αCD8^D^ nanobodies) encoding the relevant nanobody sequence, followed by an LPETGG-6xH sequence. Nanobody αCD8^M^ and αCD8^D-2^ contain the same recognition domain sequence. Protein production was induced with IPTG (Thermo Fisher Scientific) and recombinant proteins were isolated from the periplasmic fraction using Ni-NTA beads (Qiagen). Following washing and subsequent elution with 50 mM Tris (pH 8), 150 mM NaCl, 500 mM imidazole, samples were purified by gel filtration chromatography on a Biosep 3000 Phenomenex column in phosphate-buffered saline (PBS). Purity of resulting nanobodies was assessed by SDS/PAGE analysis and material was concentrated using an Amicon 10 kDa MWCO filtration unit (Millipore). Nanobodies were stored at -80 °C until further use.

### Labeling of nanobodies with photoswitchable tag or fluorochromes

Maleimide dyes (maleimide-AF647, maleimide-FITC) and the photoswitchable tag (PsT) were all coupled to GGGC peptide by incubation of 1 mg (∼20 μg/ml) of the fluorescent maleimide with 175-200 μM GGGC peptide for a minimum of 2 hours at room temperature in 10-12.5 mM NaHCO_3_. Subsequently, conjugates were purified by reverse phase HPLC on a C18 column (Waters) and identity of the obtained material was confirmed by mass spectrometry. Resulting molecules were coupled to the indicated nanobodies by sortase reactions. In brief, 2.5 μM purified nanobody-LPETGG-6xH protein was incubated with 40 μM GGGC-dye and 0.4 μM hepta-(7M) mutant sortase for 2 hours at 4 °C in 50 mM Tris (pH 8) and 150 mM NaCl. Hepta-(7M) mutant sortase was produced in-house as described^37^. Unreacted nanobodies and sortase were removed by adsorption onto Ni-NTA agarose beads (Qiagen). Subsequently, the suspension was added on top of a 100 kDa cut-off filter to remove Ni-NTA agarose beads, and flow-through (containing labeled nanobody and unconjugated GGGC-dye) was concentrated. Subsequently, unconjugated GGGC-dye was removed using an Amicon 10 kDa MWCO filtration unit (Millipore), and the material was further purified using a zeba spin column (Thermo Fisher Scientific). Labeled αCD8^M^ and αCD8^D^ nanobodies were stored in PBS at -20°C. Protein concentrations were determined by spectrophotometry and individual batches of labeled VHHs were titrated for optimal usage (final concentrations ranging from 5-10 μg/ml).

### Staining of cells and tumor tissue with αCD8 nanobodies

CD8^+^ T cells were isolated from PBMCs using the CD8^+^ T cell Isolation Kit (Miltenyi Biotec) and were stored at -80 °C in 90% FCS with 10% DMSO. At day -1, cells were thawed in RPMI (Gibco) supplemented with 10% human serum (HS, Sigma), penicillin (100 U/ml, Roche), streptomycin (100 μg/ml, Roche) and benzonase (250 U/ml, Novagen). CD8^+^ T cells were cultured overnight in RPMI supplemented with 10% HS, penicillin, streptomycin and recombinant hIL-2 (60 IU/ml, Novartis). Viable human tumor tissue pieces of ∼1-2mm^3^ were thawed in prewarmed DMEM (Gibco) supplemented with 10% FCS, penicillin (100 U/ml), streptomycin (100 μg/ml), sodium pyruvate (1 mM, Sigma), MEM non-essential amino acids (1x, Sigma) and GlutaMax (2 mM, Gibco). Tumor tissue was subsequently washed three times by thoroughly submerging and shaking the tissue pieces in fresh prewarmed medium. Where indicated, CD8^+^ T cells, PBMCs and tumor tissue were stained with αCD8^D-1^-PsT and αCD8^D-2^-FITC in PBS supplemented with 0.5% BSA (Sigma) and EDTA (2 mM, Life Technologies) for 30 minutes at 4 °C while gently shaking. Cells and tumor fragments were washed with PBS supplemented with 0.5% BSA and EDTA, and taken up in RPMI with penicillin and streptomycin and 10% HS (CD8^+^ T cells and PBMCs), or PBS with 0.5% BSA and EDTA (tumor tissue) for imaging by confocal microscopy.

### Analysis of nanobody binding stability

CD8^+^ T cells were stained with either αCD8^M^, αCD8^D-1^ or αCD8^D-2^ (FITC or AF647 labeled) as indicated (Fig. 2B and S2) in PBS supplemented with 0.5% BSA and EDTA (2 mM) for 30 minutes at 4 °C. After three washes with PBS containing 0.5% BSA and EDTA, the stained CD8^+^ T cells were mixed as indicated, followed by a 30 minutes incubation at 37 °C in RPMI with penicillin, streptomycin and 10% HS. Afterwards, cells were resuspended in PBS with 0.5% BSA and EDTA (2 mM) buffer and analyzed by flow cytometry.

### Viral transduction of tumor cells and T cells

OVCAR5 cells were cultured in IMDM medium (Gibco) supplemented with 10% FCS, penicillin (100 U/ml), streptomycin (100 μg/ml) and GlutaMax (1×). Ag^+^ GFP^+^ OVCAR5 cells and Ag^-^ Katushka^+^ OVCAR5 cells were generated by retroviral transduction with a pMX-CDK4_R>L_-GFP vector and lentiviral transduction with a pCDH-CMV-katushka-p2A-Cre^ERT2^ vector, respectively, as described previously^38^. CD8^+^ T cells were cultured in a 1:1 mix of Aim 5 (Gibco) and RPMI, supplemented with 10% HS, penicillin (100 U/ml), streptomycin (100 μg/ml) and recombinant hIL-2 (60 IU/ml). CD8^+^ T cells were retrovirally transduced with a pMP71 vector encoding the CDK4_R>L_-specific TCR (clone 17, NKI12)^39^ as described previously^40^. After transduction, T cells were expanded for two weeks using a rapid expansion protocol^41^, and cells were stored in liquid nitrogen in 90% FCS and 10% DMSO until further use.

### Tumor cell -T cell cocultures

3-5 days before coculture, tumor cells were plated in small droplets (±5,000 cells per 5ul droplet) on polymer or glass bottom 8 well μ-slides (Ibidi) in IMDM medium supplemented with 10% FCS, penicillin (100 U/ml), streptomycin (100 μg/ml) and GlutaMax (1×). Ag^+^ and Ag^-^ tumor cells were plated as indicated per experiment. Following tumor cell adherence, remaining non-adherent cells were removed by washing and cells were cultured in IMDM with 10% FCS, penicillin, streptomycin and GlutaMax. Subsequently, αCD8^D-1^-PsT and αCD8^D-2^-FITC stained CDK4_R>L_ TCR^+^ CD8^+^ T cells were added, and cells were cultured for 4 hours at 37 °C in the climate chamber of a Leica SP8 Confocal system (Leica Microsystems) microscope, as discussed below.

### Confocal microscopy imaging and local uncaging

All images were acquired using an inverted Leica SP8 Confocal system equipped with 4 tunable hybrid detectors, visible lasers (405 nm Argon, DPSS 561 nm, and HeNe 633 nm), and an Insight X3 multi-photon laser (Spectra Physics). All images were collected at 12 bit and acquired with a 25x water immersion objective with a free working distance of 2.40 mm (HC FLUOTAR L 25x/0.95 W VISIR 0.17). Fluorophores were excited as follows: FITC and GFP at 488 nm, AF594 and Katushka at 561 nm, and AF647 at 633 nm. FITC and GFP signals were collected between 510-590 nm, AF594 signal was collected between 610-650 nm, Katushka signal was collected between 620-720 nm, and AF647 signal was collected between 680-750 nm. CD8^+^ T cells or PBMCs were seeded in a Micro-Insert 4 well u-Dish (Ibidi), and placed onto the microscope with a climate chamber adjusted to 37 °C. Similarly, tumor cell – T cell cocultures were imaged and incubated in 8 well μ-slides in the climate chamber at 37 °C. Overview scans of the entire well were acquired. Tumor tissue was placed in between two cover slips (Duran), and was kept ice-cold using custom-made cool packs during image acquisition. Both overview scans and three-dimensional tile scans of the entire tumor fragments (with 1 µm Z-step size) were acquired. To uncage the αCD8^D-1^-PsT in defined areas, a population of cells was selected by drawing a region of interest (ROI). For each defined ROI, a Z-stack was made with step sizes of 1 μm. Unless indicated otherwise, uncaging was performed using the 405nm laser line at 15% power (equivalent to 865 μW/mm^2^), 600Hz, 25x magnification, and 1024 × 1024 pixels with a pixel dwell time of 600 nanoseconds.

### Harvest and dissociation of cells and tumor fragments

CD8^+^ T cells and PBMCs were harvested by gently pipetting the cells up and down three times, followed by thorough rinsing of the wells with cold PBS supplemented with 0.5% BSA and EDTA (2 mM). In case of tumor cell -T cell cocultures, non-adherent cells were harvested by gently resuspending them. Adherent cells were subsequently trypsinized with PBS supplemented with trypsin-EDTA (1x, Thermo Fisher Scientific) for 4 minutes at 37 °C to ensure harvest of a single cell suspension of tumor cells and T cells, and harvested cell fractions were pooled for subsequent staining steps. Tumor tissue was dissociated by incubation with collagenase IV (1 mg/ml, Sigma-Aldrich) and pulmozyme (12.5 μg/ml, Roche) in RPMI for 20 minutes at 37°C. After dissociation, cell suspensions were filtered through a 35 μm cell strainer (Falcon tube with cell strainer cap, Corning) and washed with cold PBS supplemented with 0.5% BSA and EDTA (2 mM).

### αFLAG staining of cell suspensions after uncaging

Single cell suspensions were stained in cold PBS with 0.5% BSA and EDTA (2 mM) either containing live-dead fixable near-IR dead cell stain (IR-dye, Thermo Fisher Scientific), αCD3-BV711 (UCHT1, BD Biosciences) and αFLAG-AF647 (D6W5B, Cell Signaling Technology), polyclonal αFLAG-AF647 (Cell Signaling Technology), or αFLAG -BV421 (L5, Biolegend) for 30 minutes at 4°C, or, when indicated, with αCD3-BV711 and unlabeled αFLAG antibody (D6W5B, Cell Signaling Technology) for 20 minutes at 4 °C, followed by addition of αRabbit-IgG-BV421 antibody (BD Biosciences) and IR-dye for an additional 15 minutes at 4°C. In tumor – T cell coculture experiments, cells were also stained with αCD69-PeCy7 (H57-597, Biolegend). Following staining, cells were washed three times and resuspended in cold PBS with 0.5% BSA and EDTA (2 mM) for flow cytometry.

### Single cell sorting of CD8^+^ T cells

CD8^+^ T cells were single cell sorted based on the following gating strategy. Forward and sideward scatter were used to exclude doublets and to distinguish CD8^+^ T cells from tumor cells. Viable CD8^+^ T cells were identified by expression of CD3, CD8 (as reflected by staining with αCD8^D-1^-PsT and αCD8^D-2^-FITC), and low IR-dye signal. In addition, CD69 expression was measured. For each sample, uncaged CD8^+^ T cells (AF594^low^ αFLAG^high^) and total CD8^+^ T cells were sorted using index sorting into 384-well plates containing 2 μl of lysis solution with barcoded poly(T) reverse-transcription (RT) primers (IDT, Li et al.^42^) per well. Four wells were left empty in each 384 well plate to be used as background controls in single cell sequencing. Following cell sorting, plates were briefly centrifuged, snap frozen on dry ice, and stored at -80°C.

### Single cell library preparation

Single cell libraries were prepared as described previously using the Massively Parallel Single-Cell RNA-seq method (MARS-seq)^32^. In brief, upon single cell sorting and cell lysis in 384 well capture plates, mRNA was barcoded and converted into cDNA. cDNA was pooled using an automated pipeline and the pooled sample was linearly amplified by T7 *in vitro* transcription. Resulting RNA was fragmented and converted into a sequencing-ready library by tagging the samples with pool barcodes and Illumina sequences during ligation, reverse transcription, and PCR. For each pool of cells, both library quality and library concentration were assessed.

### MARS seq data processing

Sequencing of scRNA-seq libraries that were pooled at equimolar concentration was done on a NextSeq 500 (Illumina) with a median sequencing depth of ∼40,000 reads per cell. Sequences were mapped to the human genome (hg19), demultiplexed and filtered as described in Jaitin et al.^32^, with the modifications reported in Li et al.^42^.

### Metacell modeling and analysis

For modeling of scRNAseq data, we used the MetaCell package^33^, using a similar strategy as described in Li et al.^42^. In brief, sets of mitochondrial genes, immunoglobulin genes, ribosomal protein genes and long noncoding RNA genes (Table S2) were removed. Cells with less than 500 UMIs were filtered out, as well as cells with a fraction of mitochondrial gene expression that exceeded 0.6. Feature genes with a Tvm=0.08 and 100 total UMIs minimal were selected. Gene features that were associated with lateral processes, such as cell cycle, type I IFN response, or stress (adapted from Li et al.^42^, Table S3) were excluded from metacell formation. Metacell generation was performed on 9,237 cells using 444 genes that passed the filtering steps. K=100 and 500 bootstrap iteration steps were done and heterogeneous metacells were split. The metacell confusion matrix was used to annotate groups of metacells that showed similar expression profiles. Three main cell groups were classified based on the expression of marker genes, including activated T cells (expressing amongst others CD8B, TNFRSF9 and CRTAM), non-activated T cells (expressing CD8B and GZMA, but lacking expression of activation markers) and tumor cells (lacking CD3D and CD8B, expressing BASP1). Tumor cells were excluded from further analysis where indicated. Supervised analysis of cell states was performed as described in the main text.

### Cell state analysis

To annotate single cells as caged or uncaged, mean fluorescence intensity values of the αCD8^D-1^-PsT and αFLAG signals per cell were used and defined as above cut-off (AF594^low^ αFLAG^high^; uncaged) or below cut-off (AF594^high^ αFLAG^low^; caged). To analyze the states of CD8^+^ T cells with different levels of uncaging, the total uncaged population was divided into five bins containing equal numbers of cells (before exclusion of tumor cells), based on their distance from the cut-off line between the uncaged and caged populations, with bin 1 containing cells with the highest level of uncaging, and bin 5 containing cells that were closest to the cut-off. For subsequent analysis of T cell states per bin, tumor cells were excluded.

The most variable genes within the dataset were defined based on their variance over all cells divided by the mean. To generate gene modules that were associated with the expression of the most variable genes, we identified the top 30 genes that correlated to one of the indicated anchor genes, IFNG, CCL4 and CXCL8, using a linear correlation of the log fold change of the expression value of a gene in each metacell over the median expression value over all metacells. Genes that were part of the cell cycle, type I IFN response, or stress modules were excluded from this analysis. The expression of gene signatures (“signature score”) for both the stress signature and the IFNG, CCL4, and CXCL8 gene modules was plotted as the fraction of signature-related UMIs of total UMIs per cell.

### Data and code availability

The processed single cell RNA sequencing data will be deposited in the NCBI GEO database. The scripts that are used for the analyses in this study will be available upon request.

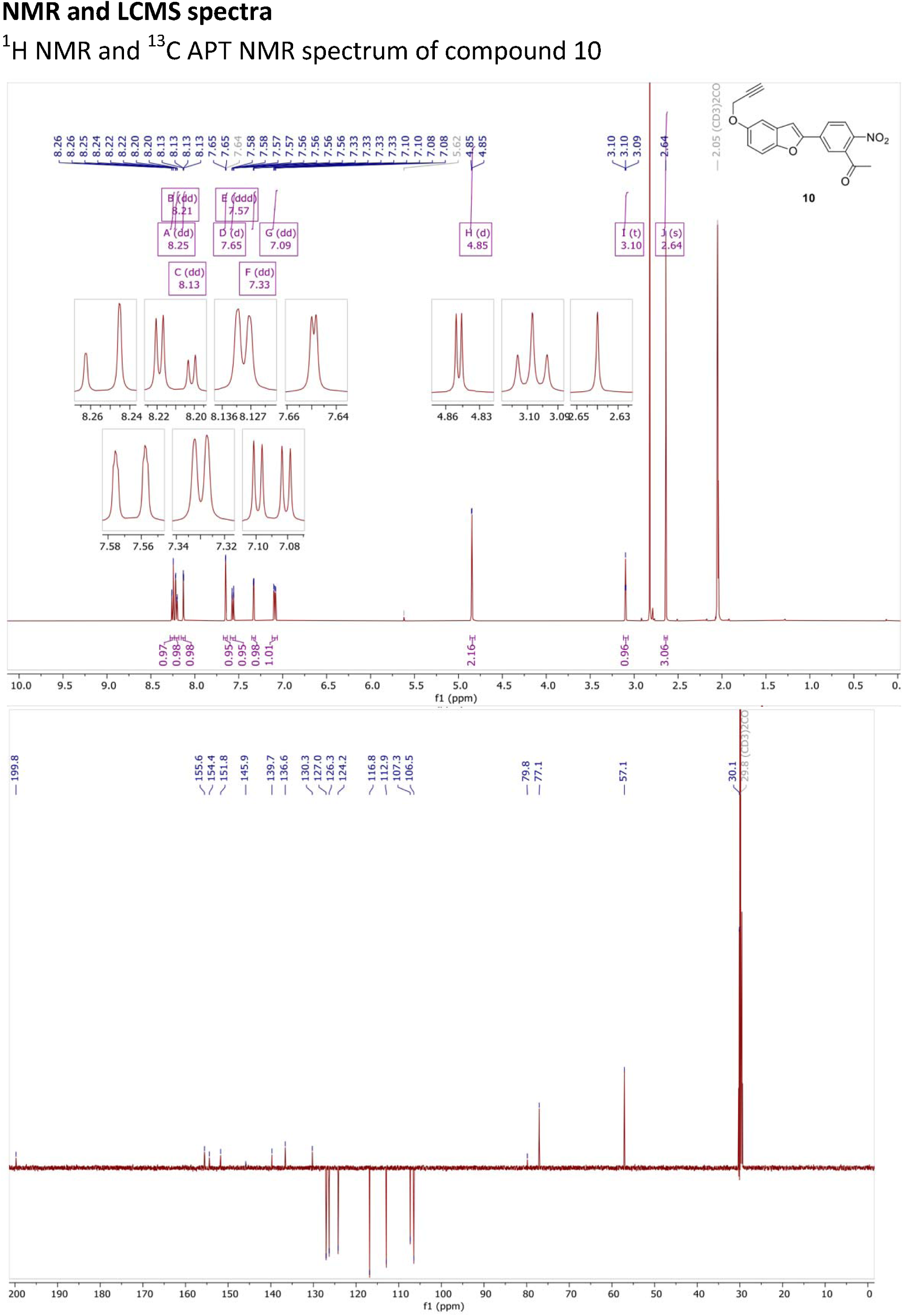

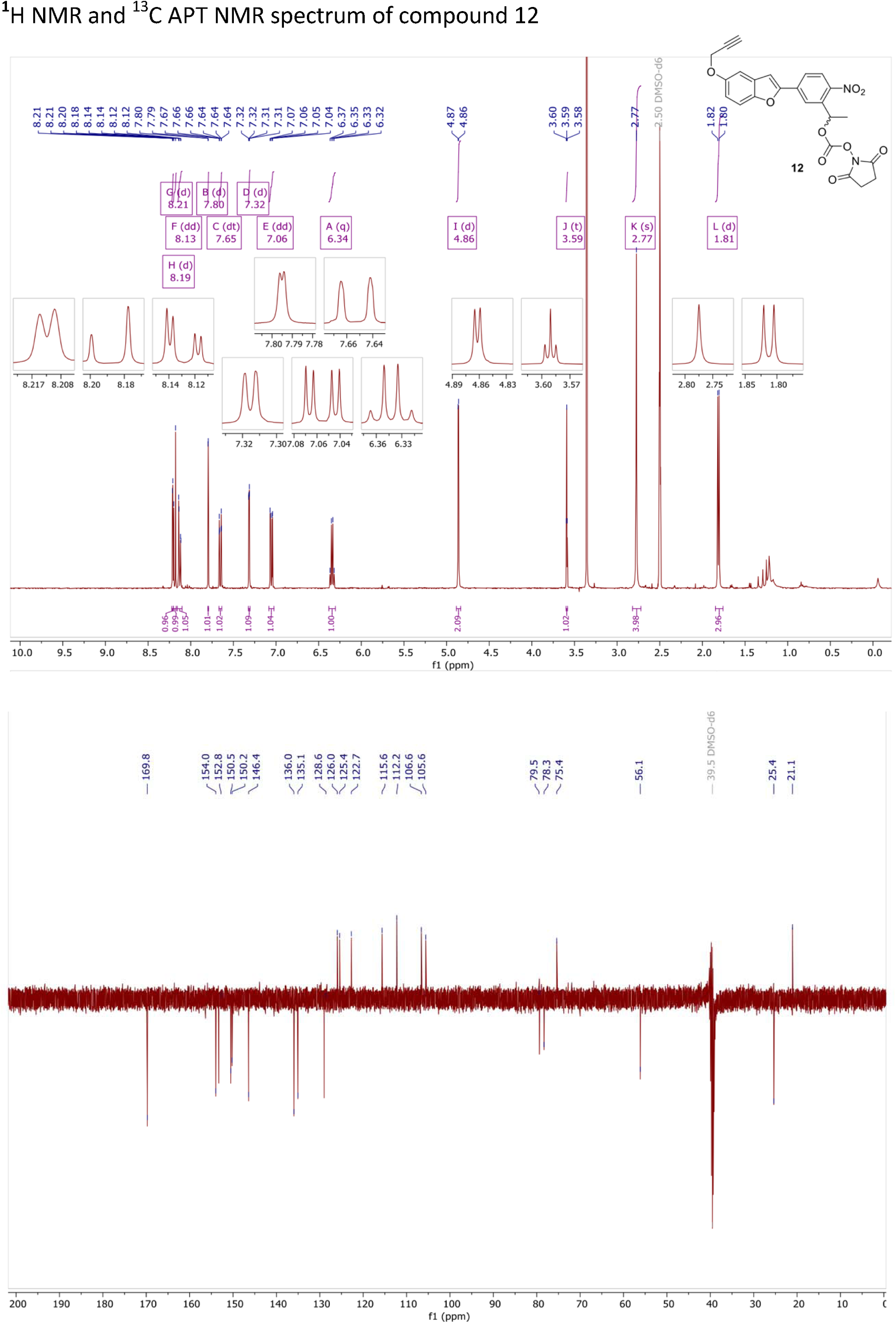

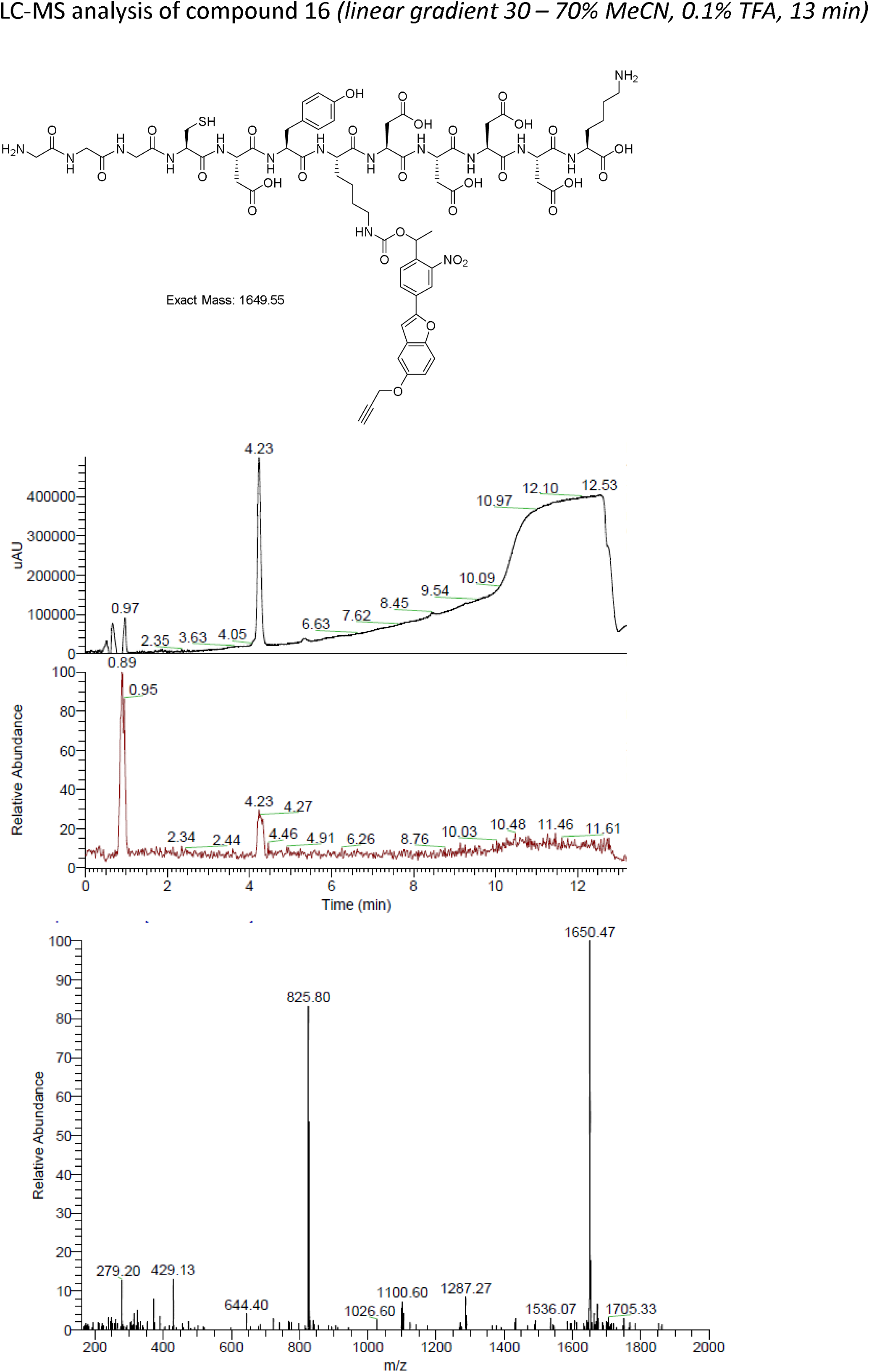

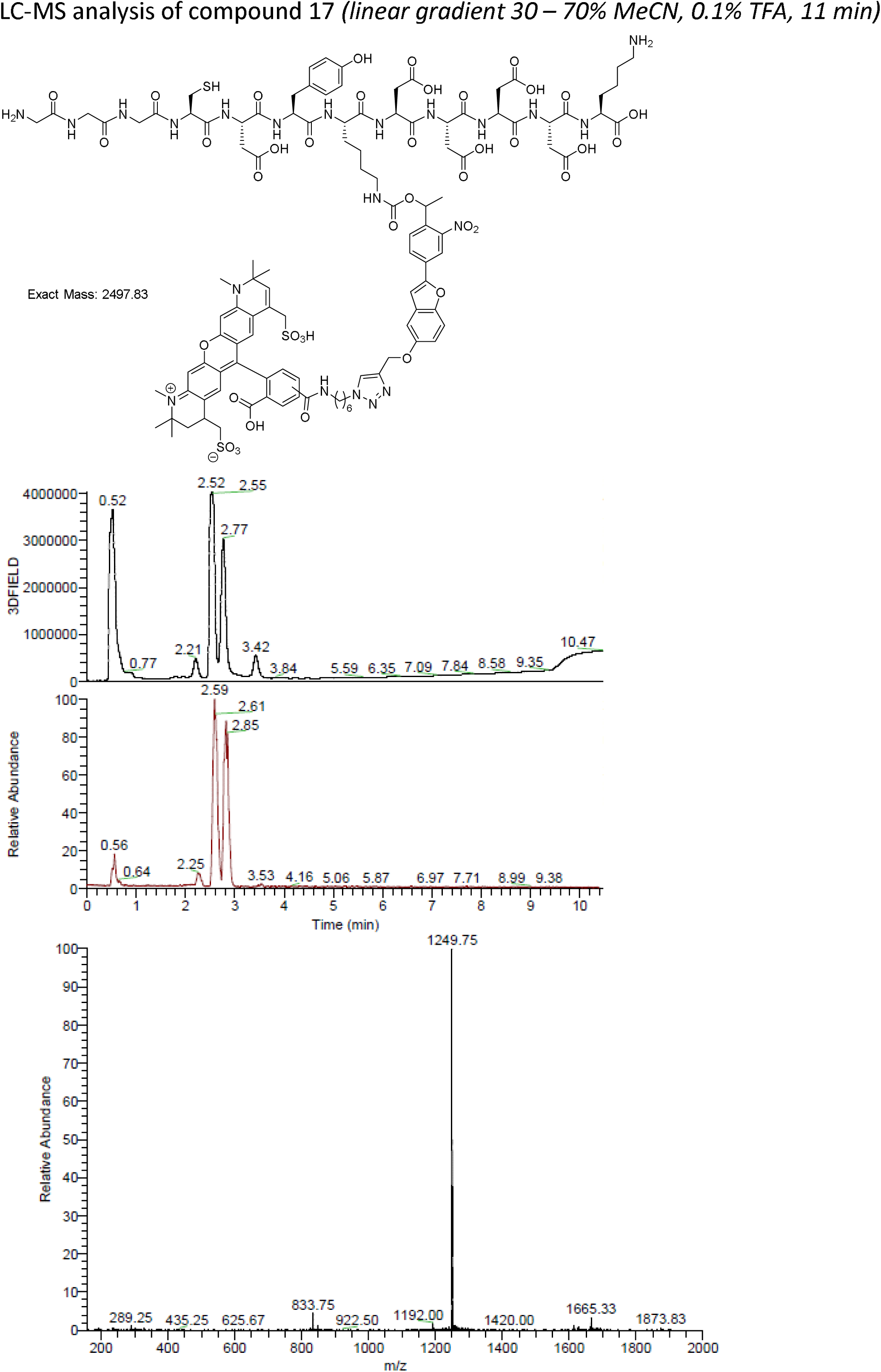

## Notes

### Competing Interest Statement

The authors have declared no competing interest.

